# Investigating Histone Lysine Acetylation and Acetyl-methylation in Mouse Brain using Enhanced Trimethylacetic Anhydride Derivatization Protocol

**DOI:** 10.64898/2026.10.08.757546

**Authors:** Hassan Hijazi, Hisham Altoufaily, Salem Al Siblani, Amandine Barbaroux, Noémie Penaud, Karine Merienne, Sabine Brugière, Delphine Pflieger

## Abstract

Histone post-translational modifications (PTMs) are key regulators of gene expression, and mass spectrometry (MS) provides a powerful platform for their characterization and quantification. However, conventional histone sample preparation using derivatization of non-modified lysines by propionic anhydride (PA) presents analytical limitations, including non-detection of short hydrophilic peptides, incomplete chromatographic separation of positional isomers, and masking of endogenous lysine propionylation and butyrylation. Here, we optimized a trimethylacetic anhydride (TMA)-based derivatization workflow to improve histone PTM analysis. Compared with PA derivatization, TMA enhanced the detection of acetylated peptides, particularly short and hydrophilic species, although peptide-dependent ionization biases affected relative peptidoform quantification. We further evaluated the suitability of TMA derivatization for investigating endogenous lysine propionylation and butyrylation. Unexpectedly, we identified lysine acetyl-methylation (Kacme) at seven sites on histone H3 and validated these assignments by targeted MS using synthetic reference peptides. To investigate the metabolic dynamics of Kacme, we performed *in vivo* isotope tracing in mice administered ^13^C-labeled glucose. Unlike canonical lysine acetylation, Kacme exhibited minimal incorporation of glucose-derived acetyl groups over 2h, suggesting substantially slower turnover of its acetyl moiety. Collectively, these findings highlight the analytical advantages and quantitative limitations of TMA-based histone derivatization, while revealing Kacme as a metabolically distinct histone modification and expanding the scope of MS-based histone PTM analysis.

## INTRODUCTION

Histone post-translational modifications (hPTMs) play central roles in chromatin biology by regulating DNA-templated processes, including replication, transcription, and damage repair. The landscape of hPTMs has grown increasingly diverse with the discovery of non-canonical modifications, including lysine lactylation^1,2^, acetyl-methylation (Kacme)^3^, pyruvylation^4^, glyceroylation^5^, and glutamine monoaminylation^6^. Mass spectrometry coupled to liquid chromatography (LC-MS) is an indispensable technique to profile the hPTM landscape with unparalleled throughput, sensitivity and quantitative accuracy. However, preparing histones for bottom-up MS analysis means specific prerequisites to address their sequence particularities^7^.

Histone proteins are indeed very rich in lysine (K) and arginine (R) residues. In the presence of trypsin, which cuts C-terminally of K and R, histones undergo excessive proteolysis, resulting in numerous small peptides (<6 residues) that are not amenable to LC-MS analysis. To circumvent this issue, derivatization reagents are used to add a chemical moiety to lysines that are unmodified (Kun) or monomethylated (Kme1, leading to Kme1pr). This restricts trypsin cleavage to only after R residues, thus producing longer and more hydrophobic peptides, which are better separated by reversed-phase LC. Additionally, this treatment constantly generates the same peptide pool regardless of the endogenous histone modification state, allowing the estimation of the relative abundances of variably modified peptidoforms with the same sequence from their respective MS signals.

Propionic anhydride (PA) is widely used in the histone proteomics community to modify Kun and Kme1 residues. Yet, PA does not confer sufficient hydrophobicity to detect short and hydrophilic peptides such as di- and trimethylated TK_4_QTAR from histone H3. Furthermore, this processing fails to provide complete LC separation of positional isomers, such as the acetylated N-terminal (4-17) of H4 which contains four lysines. Nuanced data processing is then necessary to accurately quantify these co-eluting isobaric species. Besides, PA derivatization inherently precludes the study of endogenous lysine propionylation (Kpr), as well as butyrylation (Kbu) and the more sparsely described acetyl-methylation (Kacme) that are isobaric with methylation plus propionylation (Kme1pr).

To mitigate these issues, a protocol using an alternative anhydride, trimethylacetic anhydride (TMA; +84.0575 u), has been described^8^. Lysine TMAylation provides enhanced separation of acetylated H4 (4-17) positional isomers, allowing their straightforward MS1-level quantification. However, the originally described protocol comprises two derivatization steps, one containing overnight incubation and the second using a short treatment in microwaves^8^, making it significantly lengthier than the standard PA protocol^9,10^.

In this work, we optimized a TMA derivatization protocol akin to the widely used PA protocol, facilitating its seamless adoption. Next, we evaluated the extent to which TMA provides similar estimations of relative peptidoform abundances compared to PA. Furthermore, we investigated the possible identification of endogenous lysine propionylation and butyrylation in raw MS data. Interestingly, close inspection of MS/MS spectra acquired on mouse brain histones tentatively matched to propionylated sequences led us to the assumption of peptides actually modified with Kacme. Comparison to synthetic peptides bearing this dual modification via targeted MS analysis allowed validating the identification of Kacme-containing peptides, exclusively on histone H3. Finally, injection of ^13^C-glucose into mice allowed probing the turnover rate of acetylation within this dual modification relative to standalone acetylation.

## RESULTS

### Simplification of the TMAylation protocol

The original TMAylation protocol requires multiple rounds of chemical derivatization, including an overnight incubation at room temperature followed by microwave-assisted reactions^8^. To align this with the PA-derivatization protocol routinely used in histone proteomics workflows^9–12^, we developed an optimized version of the TMAylation protocol.

Firstly, to obtain efficient labeling, the concentration of TMA was substantially increased relative to that used for PA (see Methods section), thereby matching the specific TMA-to-histone ratio described in ^8^. When applying this modified protocol, we observed notable undesired side reactions on Ser (S) and Thr (T) residues, similar to what was previously reported during derivatization with PA^13^ (**Suppl. Figure 1**). This side product, called overlabeling, was efficiently reversed using a subsequent one-hour basic treatment^13^ (**Suppl. Figure 1**).

Secondly, we optimized the LC gradient typically utilized for propionylated histone peptides to account for the greater hydrophobicity imparted by the trimethylacetic groups. As detailed in the Methods section, increasing the slope of the acetonitrile gradient allowed for improved utilization of the earlier elution window (**Suppl. Figure 2**). Under these optimized separation conditions, we could verify the excellent separation of acetylated positional isomers, particularly those of the H4 (4-17) peptide, which contains four lysine residues. This was described as the primary advantage of TMA derivatization chemistry^8^. Notably, utilizing an Aurora C18 column yielded even greater separation of these N-terminal H4 positional isomers (**Suppl. Figure 2**).

Thirdly, we evaluated the relative abundances of fully TMAylated (desired), underlabeled (incompletely derivatized), and overlabeled peptides within our LC-MS/MS data, benchmarking our results against those obtained using the original TMA protocol^8^. As shown in **Suppl. Figure 3** for peptides containing from one to three lysine residues, our protocol provided a higher yield of fully labeled peptidoforms. Of note, only the subset of desired peptides is usually utilized for downstream quantitative analysis, assuming almost complete labeling.

Lastly, we evaluated the labeling efficiency of endogenously monomethylated peptides across both protocols, focusing on those from the histone H3 N-terminus that were consistently and confidently identified (**Suppl. Figure 4**). Kuchaříková *et al*. attributed the low labeling efficiency of these species to steric hindrance caused by the bulkiness of the trimethylacetyl group^8^. Notably, we found that the TMA-me1 to me1 ratio for these peptides was substantially higher using our optimized protocol than with the original method, which exhibited poor labeling of monomethylated lysines (**Suppl. Figure 4**).

Overall, we developed an adapted TMAylation workflow that aligns seamlessly with standard histone preparation pipelines. Our optimized protocol demonstrated significantly higher derivatization efficiency at non-modified and monomethylated lysines, and minimized overlabeling at S/T residues compared to the original method.

### TMAylation distorts the estimation of relative abundances of variably modified peptides, to favor in particular acetylated sequences

Endogenous PTMs and chemical modifications may significantly impact the ionization efficiency of a given peptide^14,15^. This is an aspect to bear in mind when analyzing histones, because the relative stoichiometry of each modified form of a proteolytic peptide is commonly estimated from the MS1 signal of a given peptidoform, divided by the sum of the MS1 signals detected for all variably modified forms of same sequence. This calculation assumes that chemical modification of lysines resembles naturally occurring PTMs, which are also considered to have a comparable effect on ionization efficiency relative to their unmodified counterparts, even though this constitutes a known approximation^15^. While this assumption may be reasonable for propionyl due to its structural similarity to endogenous acetylation, the question is worth asking how much the very bulky and hydrophobic trimethylacetyl structure alters estimated stoichiometries relative to conventional PA-based measurements. This technical question was previously investigated for various other anhydrides by Simone Sidoli *et al*.^16^.

To address this question, mouse testis histones were prepared in technical triplicate and processed using either TMA or PA. Some histone peptides yielded highly consistent estimations of relative stoichiometries, as observed for the H3 (18-26) peptide (K_18_QLATK_23_AAR), even though monomethylation at either H3K18 and H3K23 was uniquely identified with PA derivatization (**Figure 1**). In contrast, highly divergent results were obtained for peptide sequences harboring complex combinations of acetylation and variable methylation states. Indeed, the H3 (9-17) peptide (K_9_STGGK_14_APR) appeared di- and-trimethylated at H3K9 under both derivatization conditions. Nonetheless, a strikingly elevated MS signal was observed with TMA for the co-occurring H3K9me2K14ac and H3K9me3K14ac peptidoforms, whereas these specific PTM combinations were quantified well below 1% using the conventional PA workflow (**Figure 1**).

**Figure 1.**
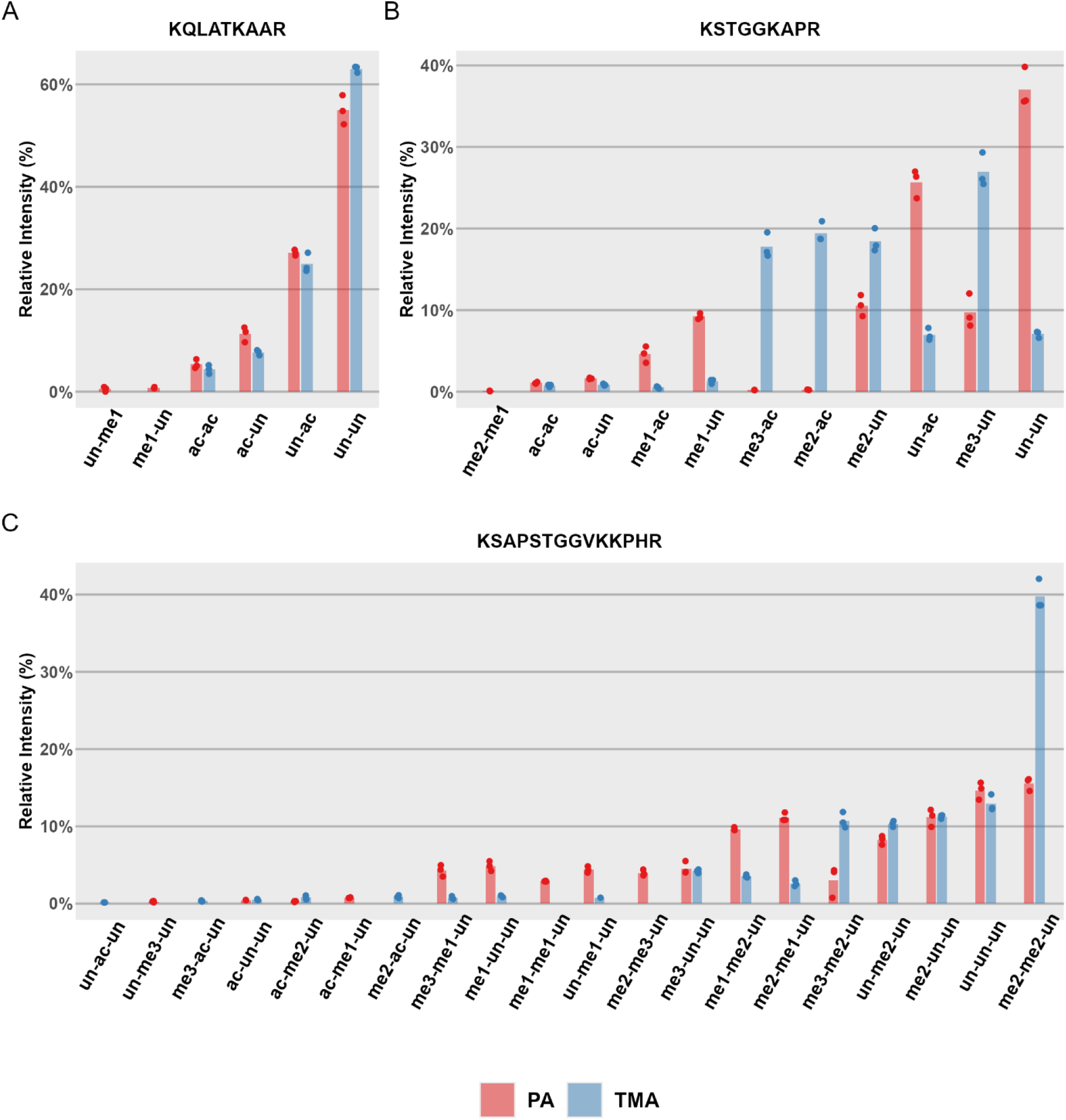
A comparison of the relative abundances of histone H3 N-terminal peptides estimated when histones are derivatized with TMA or PA. The sequences are: A) K_9_STGGKAPR_17_, B) K_18_QLATKAAR_26_, and C) H3.3 K_27_SAPSTGGVKKPHR_40_. The height of the bar represents the mean relative abundance estimated from three technical replicates represented as points. On the x-axis are indicated the modification combinations at Lys residues contained in the considered peptides. All the peptides are N-terminally labeled. “un” refers to a Lys that is chemically TMAylated or propionylated.

Another notable example was found within the H3 (27-40) sequence. In particular, the H3.3 variant showed fewer peptidoforms containing H3K36me1 when using TMA. Conversely, this reagent substantially enhanced the detection of peptides acetylated at K27 or K36, wherein the remaining lysine residues were either unmodified or methylated (**Suppl. figure 5**). The scope of identified acetylated sequences was further expanded to include specific sites at H3K4, H3K36, H3K56, H3K64, H3K79, H4K31, H4K59, H4K77. This benefit also extended to numerous lysine sites on H1, H2A, H2A.J, H2A.X, H2A.Z, TSH2A, macroH2A and H2B, facilitating the robust characterization of short and hydrophilic peptides, including K_36_GNYSER, AK_13_AK_15_TR and AK_13_AK_18_SR from H2A, as well as GK_5_TGGK_9_AR from H2A.X (**Suppl. figure 5**).

Overall, TMA derivatization introduces a bias in peptide ionization efficiency compared to PA, although the effect is not uniform among all sequences. Under our conditions, this modification notably favors the ionization of acetylated species, which we detect on many short peptides from various histone proteins.

### TMA allows detection and characterization of acetyl-methyl lysine modification

In addition to improving chromatographic separation of positional isomers and enhancing the ionization efficiency of acetylated peptides, the TMA workflow might enable the investigation of endogenous lysine acylations that are inaccessible when using conventional PA-based derivatization strategies. In particular, TMA theoretically allows the analysis of endogenous propionylations and butyrylations. We ran a database search on data acquired on histone samples extracted from mouse brain hemispheres, while specifying these two PTMs as variable modifications. Interestingly, while no butyrylated peptide was identified, a substantial number of experimental MS/MS spectra (26 spectra with Mascot scores >25) were assigned to propionylated sequences, specifically derived from histone H3. However, closer examination of the MS/MS spectra revealed the lack of cyclic immonium (CycIm) ion characteristic of Kpr at m/z 140.1075, which is expected to be particularly intense when this modification is present on the lysine in first position of the peptide^10^. They instead contained an abundant fragment ion at m/z 98.097, which is usually produced by monomethylated lysines^10^. This observation prompted us to consider other modifications strictly isobaric to propionylation, and including a methyl group.

Previous reports described that lysine residues can simultaneously carry acetylation and monomethylation on the ε-amino group, defining the dual modification termed acme which exactly matches the mass of propionylation^3,17^. We have extensively observed that the Kme1pr-peptides generate the CycIm ion characteristic of Kme1 (**Suppl. Figure 6**). The fragmentation patterns observed on histone peptides could then be attributed to the dual modification acme on lysine (Kacme). To validate this hypothesis, we performed parallel reaction monitoring analysis on mouse striatal histones targeting light endogenous peptides tentatively harboring this modification, along with their synthetic heavy counterparts bearing the Kacme moiety which were spiked into the samples (**Suppl. table 2**). Fourteen endogenous peptidoforms covering seven distinct lysine residues from H3 (H3K4, K9, K18, K23, K27, K36, K79) were confidently validated: both the endogenous and the heavy-labelled counterparts displayed identical retention times and twin MS/MS fragmentation patterns, including the characteristic m/z 98.097 CycIm ion (**Figure 2**). Intriguingly, no Kacme-containing peptides were detected in histone H4 (**Figure 2**), despite a detailed study identifying this modification on the H4 N-terminal tail across multiple species and mammalian tissues, including the brain^3^.

**Figure 2.**
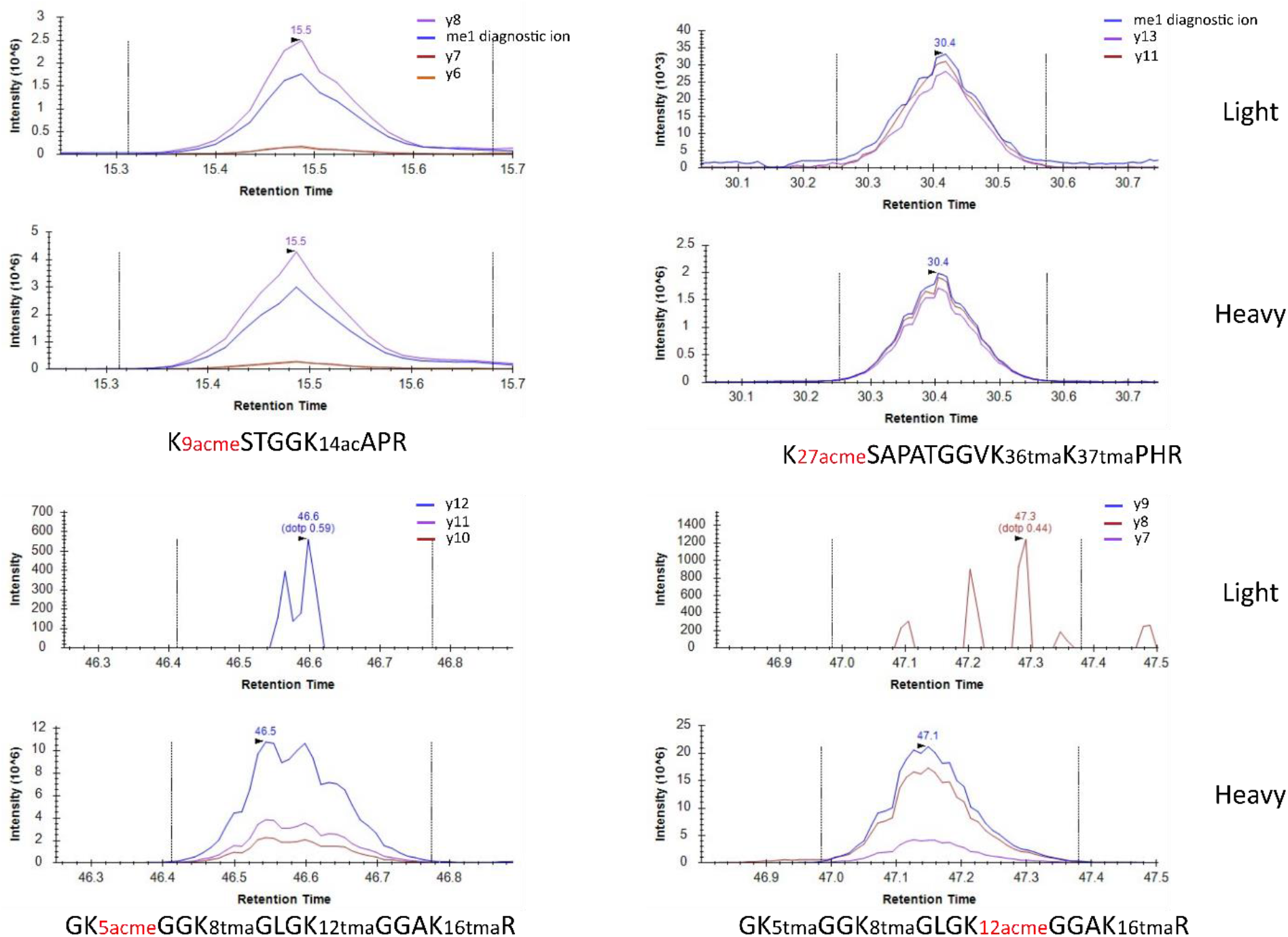
LC-MS elution profile of heavy synthetic peptides bearing Kacme with possibly corresponding endogenous peptides. Selected MS/MS transitions over the elution peaks of heavy synthetic Kacme-containing peptides and their possible light endogenous counterparts are shown. Peptides modified by H3K9acmeK14ac and H3K27acmeK36unK37un showed co-elution of the light and heavy peptides together with coherent fragment ion profiles. In contrast, no endogenous signal co-eluting with the corresponding heavy synthetic peptide was detected for peptides modified by H4K5acmeK8unK12unK16un or H4K5unK8unK12acmeK16un.

Our initial MS1-based quantification indicated that several of these Kacme-modified peptides were of comparable or higher abundance than their corresponding acetylated forms **(Table 1. A)**. This trend could be verified by manually extracting the corresponding MS1 signals (**Suppl. Fig. 7**). We next estimated the abundance of the dual modification in each peptidoform relative to the acetylated counterpart based on the integrated peak areas of MS2 fragment ions. Unexpectedly, abundance estimates differed substantially from those obtained during the initial discovery analyses based on MS1 signals **(Table 1. B)**. For example, the peptide H3K9acmeK14un which initially appeared 4 times more abundant that the corresponding acetylated form, was consistently detected at markedly lower abundance during subsequent PRM analysis, both when considering MS1 or MS2 quantification. Because the analytical conditions were comparable, we investigated whether prolonged storage at - 80°C affected stability of Kacme-modified histones. However, analysis of histones extracted from freshly prepared tissue failed to reproduce the high signal intensity initially observed **(Table 1. C)**. Although the underlying cause of variable relative abundance compared to the acetylated counterparts remains unclear, Kacme-containing peptides were reproducibly detected across independent experiments. In all, we have confidently identified the dual mark Kacme at 7 lysine sites of histone H3, with relative abundances compared to the canonical acetylation mark reaching at least 18-25% at residues H3K9 and H3.3K27.

**Table 1.** Comparison of Kacme-peptide abundances relative to the corresponding Kac peptidoforms. Values represent the mean of ratios over replicates of the signal of each Kacme-containing peptidoform to that of its corresponding acetylated form. ***A***, Initial discovery analysis based on MS1 signals (N=4 biological replicates); ***B***, Targeted MS2-based quantification performed on stored samples (N=3 technical replicates); ***C***, Targeted MS2-based quantification performed on freshly prepared brain samples (N=3 biological replicates). N/A, ratio not available. Md: modified; designates the histone H3 lysine where the modifications acme and ac are compared.

| Ratio<br>acme/ac | K4md | K9mdK14un | K9mdK14ac | K18mdK23un | K18mdK23ac | K18unK23md | H3.3K27md | H3K27me3K36md | K79md |
| --- | --- | --- | --- | --- | --- | --- | --- | --- | --- |
| A | 6.44 | 4.28 | 1.39 | N/A | N/A | N/A | N/A | N/A | 11.96 |
| B | 2.12 | 0.18 | 0.08 | 0.0047 | 0.0014 | 0.0001 | 0.39 | 0.3 | 0.28 |
| C | 0.79 | 0.28 | 0.15 | 0.0118 | 0.0011 | 0.0006 | 0.25 | 0.09 | 0.11 |

Finally, we investigated the dynamics of the acetylation moiety within acme and compared it with the canonical acetylation mark. For this purpose, we extracted histones from brain hemispheres belonging to mice pre-injected with ^13^C-labelled glucose (see **Material and Methods**), and sacrificed after 30, 60 or 120 min. As expected, the acetylated peptidoforms containing H3K9, H3K18 and H3K27 exhibited progressive incorporation of the ^13^C-labelled acetyl group during the first hour before it started decreasing. In contrast, little or no incorporation of heavy acetyl label was detected for the Kacme-containing peptidoforms, indicating that the acetyl moiety within the dual modification remained largely unchanged over 2 hours **(Figure 3)**.

**Figure 3.**
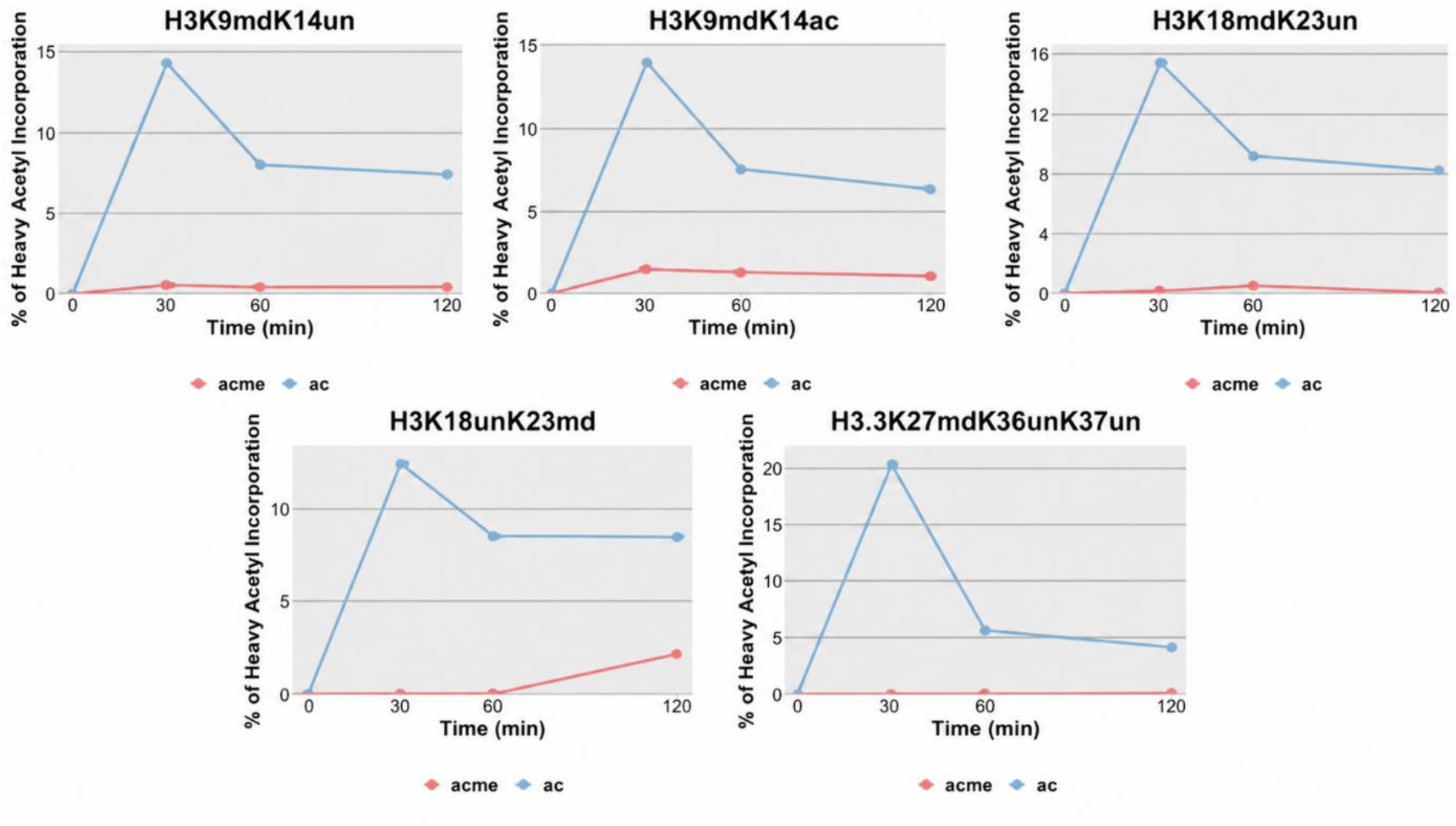
MS2-based comparison of ^13^C-acetyl incorporation into conventional acetylated and Kacme-containing H3 peptidoforms after intraperitoneal injection of ^13^C-glucose in mice. Percentage of heavy acetyl incorporation was followed at 0, 30, 60 and 120 min after ^13^C-glucose administration for the indicated H3 peptidoforms. Md: modified; designates the histone H3 lysine where the modifications acme and ac are compared.

## DISCUSSION

### Valeric anhydride as an isomer of TMA previously considered for histone derivatization

Sidoli *et al*. had tested various lysine-modifying chemistries for derivatizing histones^16^. PA was finally selected as the best reagent due to the production of high quality MS2 spectra and reasonable deviation of MS1-based estimated abundances from standard synthetic peptides of known concentration. Among the tested compounds was a structural isomer of TMA, valeric anhydride which has linear unbranched acyl chains (**Suppl. Fig. 8**). Though this derivatization agent led to an advantageous shift in RT and peptides exhibited good quality MS2 spectra, it induced significant deviations of abundances compared to known-ground-truth standards^16^. Additionally, fewer peptides were detected compared to what PA afforded. These missing identifications mainly included doubly and triply methylated H3 (9-17) peptides which were well detected with our optimized TMA protocol (**Fig. 1B**). This improvement of TMA over valeric anhydride merited further investigations.

### Increased coverage of acetylation sites and detection of Kacme instead of Kpr using TMA

When comparing the list of identified peptides using PA or TMA chemistries, we observed the identification of many more acetylated peptides with TMA, including very short sequences generated by multiple H2A variants. This advantage may be beneficial when one wants to study the dynamics of acetylation, such as in tracing experiments aiming at following the flux of heavy labelled metabolites (e.g. ^13^C-acetate, ^13^C-glucose) into histone acetylation. Besides, unlike when derivatizing histones with PA, endogenous propionylation and butyrylation might be studied when using TMA. However, our results showed that TMA did not afford identification of butyrylated peptides, and the fragmented sequences tentatively attributed to propionylated sequences actually corresponded to other isobaric peptidoforms, namely modified by Kacme. This identity emerged from the absence, in MS/MS spectra, of the CycIm ion produced by Kpr (at m/z 140.1075) and the detection of the CycIm produced by Kme instead (at m/z 98.097). The latter fragment ion is also produced by peptides containing the dual modification Kme1pr (coming from the chemical propionylation of endogenous methylated lysines), which encouraged us to test the hypothesis of the dual Kacme modification, using heavy-labeled synthetic peptides. Its presence was confirmed to occur on all inspected lysines of histone H3, namely H3K4, H3K9, H3K18, H3K27 and H3K79, but was ruled out on all lysines of histone H4, including on its N-terminal where it was formerly functionally characterized^3^. It is interesting to note that we detected Kacme on all H3 lysines which are known to alternate between acetylation and di/trimethylation.

When analyzing successive samples, we observed discrepancies in the abundance of Kacme peptides estimated by targeted analysis compared to what was conferred from the discovery (MS1-level) data. We ruled out storage at -80 °C of the tissue as a possible cause by analyzing freshly prepared samples. Even though the relative abundance of Kacme compared to Kac remains to be ascertained in various tissues and conditions, our quantification data indicate non-negligible abundances, especially at H3K4, H3K9, H3K27 and H3K79.

Last, to assess the dynamic nature of Kacme compared to Kac, we followed the incorporation of ^13^C atoms into each acetyl moiety over 2h after intraperitoneal injection of ^13^C-glucose into mice. While acetylation at lysines of the N-terminal of H3 exhibited 13-20% heavy atom incorporation, a negligible metabolic turnover rate was observed for the acetyl moiety of Kacme. This aligns closely with biochemical findings demonstrating that Kacme is structurally resistant to enzymatic deacylation by classical Zn^2+^-dependent histone deacetylases (HDACs 1–11) and NAD^+^-dependent sirtuins (SIRT1– 7)^3,18^. Together, the extremely reduced metabolic and enzymatic dynamics of Kacme reinforce the functional model describing this hPTM as a quasi-permanent chromatin mark that safeguards open chromatin regions, sustains local DNA accessibility, and acts as a guardian of the genome to oppose heterochromatin spreading and broad chromatin compaction^3,19^.

### Only CycIm ions allowed rejecting Kpr and investigating Kacme

Our work highlights again how critical it is to visually inspect MS/MS spectra acquired on histone peptides to validate identifications of acylated sequences, including by looking for cyclic immonium (CycIm) ions in the low-mass range. The importance of CycIm ions to distinguish isobaric peptidoforms has been emphasized in previous publications^10,20^, and CycIm ions were used by Tang *et al*. to tell apart Kacme from Kpr at H3K27^17^. Indeed, the peaks at m/z 98.097, 126.091, 140.107, and 168.128 corresponding to Kme, Kac, Kpr and Ktma, respectively, are expected to be very intense when the modified lysine is in the first position of the peptide. Research reporting endogenous propionylation should always verify these respective CycIm ions before moving forward to biological conclusions. This stresses also the advantage of search engines that facilitate the utility of CycIm ions, such as Andromeda and MSFragger^21–23^, in reducing false identifications^10^. Moreover, ion mobility could potentially be used to separate Kacme- and Kpr-modified peptides, as both have different sizes and structures. Of note, ion mobility has been successfully used to resolve isobaric and isomeric PTM complexity in histones, and it may be worth incorporating it into proteomic mass spectrometry workflows as it enhances sensitivity and boosts the number of detected histone peptides^24,25^.

### Possible disadvantages of the derivatization with TMA

TMA would theoretically mask other endogenous PTMs such as isovalerylation and 2-methylbutyrylation^26^. While these modifications have not yet been described to occur on histones^27^, the presence of their corresponding acyl-CoA make them potential candidates^26^. Moreover, it is worth noting that some PTM combinations including TMA are nearly isobaric with a mass difference below 3 ppm (TMA *versus* me2 + pr; TMA+me1 *versus* me3+pr; TMA-ac *versus* bu + pr). Nonetheless, the overall strictly isobaric peptidoform space created by TMA is narrower than that created by PA. An example showing this comparison on H3.1 and H3.3 K27-R40 stretch by modeling possible PTM×sequence combinations is illustrated in **Suppl. Fig. 9**.

It is also important to note that the precursor charge state distribution is affected by the derivatization reagent. Under the conditions used here, TMA derivatization produced a higher proportion of 2+ precursor ions, whereas PA derivatization favored 3+ precursor ions (**Suppl. Fig. 10**). Both trimethylacetylation and propionylation convert the lysine ε-amino group into a neutral amide and therefore remove a potential protonation site. The precise physicochemical basis of this reagent-dependent charge-state shift remains to be established. The lower sequence coverage but higher number of peptide identifications observed with TMA may be partly related to the greater hydrophobicity introduced by the trimethylacetyl group, which can improve chromatographic separation of modified peptide forms but also reduce the recovery or detectability of larger, more hydrophobic peptides, as reported by^8,27^. When comparing several histone-MS workflows, PA derivatization produced a substantial proportion of singly charged precursor ions, and the H3(9–17) peptide showed a particularly large singly charged contribution, with approximately 60% of its signal assigned to the 1+ state^12^. Because singly charged features may be detected at the MS1 level but are usually excluded from MS/MS fragmentation, they could contribute to discrepancies between MS1-signal-based relative abundance measurements and the number of MS/MS-based peptide identifications. This possibility may be especially relevant for the H3 (9–17) peptide, for which we observed inconsistent relative abundances between PA and TMA protocols.

### Methylamine as an alternative to hydroxylamine

Histone derivatization reaction is not flawless, causing O-derivatization of S/T/Y residues. To address this issue, which compromises LC-MS analysis sensitivity and quantitation accuracy, two approaches have been devised: (i) preventive, which maintain a pH around 8 ^11^, and (ii) curative, which adds a reverse-labeling step after derivatization reaction^13^. The latter is done by applying hydroxylamine at a high pH, a step implemented in our protocol. The original TMA protocol used the preventive method by adjusting pH to be around 8 with NH_4_OH^8^. Using hydroxylamine does not fully guarantee the removal of all O-TMA with traces remaining even with prolonged incubation as shown in **Suppl. Fig 1**.

A recent study has demonstrated that using methylamine compared to hydroxylamine exhibited greater efficiency (>99%) to selectively reverse O-acyl formation after N-Hydroxysuccinimidyl (NHS) ester-mediated derivatization^29^. However, since methylamine is dissolved in methanol, careful consideration must be given to the possibility of other side reactions like artifactual methylations^30^. Methylamine is also worth testing in TMA- and PA-workflows.

### ArgC-based protocols are gaining momentum

Besides, the adoption of an ArgC-based protocol instead of trypsin is gaining traction in histone proteomics, driven by the advent of recombinant ArgC enzymes^12,28,31,32^. While conventional Arg-C preparations suffered from low specificity and inconsistent cleavage^12,33^, modern recombinant proteases such as Arg-C Ultra and Arg-C Zero operate with near-complete cleavage efficiency (>97– 99%)^28,31,32^ encouraging their immediate adoption. For long, lysine-rich peptides such as the N-terminal peptide of histone H2B, an ArgC-only workflow evades generating very hydrophobic peptides with prolonged retention times and higher risk of incomplete labeling in sequences containing 10 lysines. Because internal lysine residues remain unmodified by chemical derivatization, Arg-C peptides often exhibit higher charge states, which complicates MS/MS spectral interpretation. Conversely, post-digestion derivatization at the peptide level remains essential for retaining short, hydrophilic peptides (e.g., H3K4). The latter workflow is especially attractive for single-cell histone proteomics to streamline the protocol without sacrificing sensitivity^34^. Nonetheless, challenges with over- or under-labeling persist, highlighting the value of our optimized protocol to maximize the labeling efficiency and minimize side reactions.

## CONCLUSIONS

We established an optimized histone derivatization protocol using trimethylacetic anhydride (TMA) that mirrors the workflow of the widely adopted propionic anhydride (PA) method. Importantly, this optimized TMA protocol enhances labeling efficiency, including that of monomethylated peptides, while successfully minimizing undesired O-TMAylation on S/T residues. Furthermore, our evaluation reveals that TMAylation alters the ionization efficiency of some histone peptidoforms, leading to distorted estimation of relative peptidoform abundances compared to results provided by the PA chemistry. In particular, we found that detection of acetylated peptides, including those of very short sequence, was favored. Lastly, we explored the detection of endogenous propionylation, which is masked when using PA. We instead detected peptides of histone H3 bearing the dual Kacme modification. We investigated this underexplored modification, by showing that it is largely insensitive to glucose in the mouse brain, in stark contrast with lysine acetylation.

## MATERIALS AND METHODS

### Chemicals, Synthetic Peptides, and Tissues

Propionic anhydride (≥99%, Product No. 240311) and Trimethylacetic anhydride (≥99%, Product No. 143502), and hydroxylamine (Product No. 438227) were purchased from Sigma Aldrich. Heavy labeled synthetic peptides were purchased from JPT Peptide Technologies.

### Animals

Wild-type (WT) mice were maintained on C57BL/6J genetic background. Animal studies were conducted in accordance with French regulations (EU Directive 2010/63/UE–French Act Rural Code R 214-87 to 126). The animal facility was approved by veterinary inspectors (authorization no. E6748213) and complies with the Standards for Human Care and Use of Laboratory Animals of the Office of Laboratory Animal Welfare. All procedures were approved by local ethics committee (CREMEAS) and French Research Ministry (no. APAFIS#31527_2021051713407251v3). Mice were housed in a controlled-temperature (22°C) and -humidity (40%) room maintained on a 12 h light/dark cycle. Food and water were available ad libitum. For proteomic analyses, mice were killed by cervical dislocation and cerebral tissues were rapidly dissected, snap frozen and stored at −80 °C until further processing.

### Heavy glucose tracing experiment

Mice were fasted for 12 hours before the experiment but had free access to water. On the day of the experiment, they were injected intraperitoneally with heavy glucose (U-^13^C_6_, Cambridge Isotope Laboratories) at a concentration of 0.225 M (≈560 mg/kg body weight per injection)^35^. Thirty minutes later, a subcutaneous injection of the same solution was administered. The first injection was intended to saturate the liver with glucose and facilitate glucose perfusion to the brain following the subcutaneous injection. Brain hemispheres were collected 30, 60, and 120 minutes after the second injection. Control mice did not receive any glucose injection.

### Histone extraction

Testis histone samples were used to test and develop the TMAylation protocol, whereas mouse brain histones were used for the stable isotope tracing experiment using ^13^C-glucose. Briefly, histone proteins were extracted from mouse brain hemispheres following a protocol developed by Maile *et al*. with modifications^9^. Cell lysis and nuclei isolation were performed using NP-40 (0.3% final concentration) in nuclei isolation buffer (NIB) (15 mM Tris-HCl, pH 7.5, 60 mM KCl, 11 mM CaCl_2_, 5 mM NaCl, 5 mM MgCl_2_, 250 mM sucrose, 1 mM dithiothreitol (DTT)). Sodium butyrate, trichostatin A, and nicotinamide were added immediately before the use of NIB to achieve final concentrations of 10 mM, 3 µM, and 50 mM, respectively. The tissue samples were homogenized using a Dounce homogenizer (A and B) in the presence of 1 mL of NP-40-containing NIB. The resulting homogenate was centrifuged to isolate the nuclei. Subsequent washes were performed to remove cytoplasmic proteins by resuspending the nuclei in NIB without NP-40. Histones were then extracted by resuspending the pellet in 1 mL of a 0.2 M H_2_SO_4_ solution and rotating the tubes at 4°C for 1 hour. The supernatant was collected after centrifugation and treated with TCA to precipitate histone proteins. After centrifugation, the supernatant was discarded, and the histone pellet was washed with 0.1% HCl in acetone stored at -20°C until use, and a second with non-acidified cold acetone. This step was followed by resuspension of histones in water. SDS gel electrophoresis of an aliquot was performed to confirm the quality of histone samples.

### Histone in vitro derivatization

TMAylation was carried out similarly to *in vitro* propionylation described elsewhere but with slight modifications. TMA solution was prepared by first making an intermediate dilution in acetonitrile (ACN) due to the poor miscibility of concentrated TMA in water. From this stock solution (246.5 mM in ACN), a 71 mM TMA solution was prepared by mixing 288 µL of it with 356 µL of 200 mM TEAB, and 356 µL of ACN, resulting in 1 mL of TMA solution.

In the first derivatization step, 20 µL of TMA solution was added to dried histone samples and incubated at 37°C for 30 minutes with shaking. A second derivatization step was followed by adding another 20 µL of freshly prepared TMA solution, incubating at room temperature for 30 minutes, and drying the samples using a SpeedVac. For N-terminal peptide modification, the derivatization steps were repeated similarly. Reverse TMAylation was performed by resuspending peptides in 500 mM hydroxylamine, adding NH_4_OH to reach pH 12, and incubating for 60 minutes at room temperature with shaking. The pH was lowered using pure TFA, followed by drying in a SpeedVac for over 3 hours. Finally, peptides were resuspended in a buffer (0.1% TFA, 5% ACN), and the pH was adjusted to 2-3 for C18 desalting (BioSPE™ PurePep).

### LC-MS/MS

Histone peptides were analyzed using Vanquish Neo UHPLC System coupled to Ascend Tribrid mass spectrometer in DDA mode. Briefly, the mobile phases consisted of solvent A (water with 0.1 % FA) and solvent B (ACN with 0.08 % (v/v) FA). The TMA-adapted LC gradient was modified from the PA-adapted gradient to accommodate more hydrophobic peptides labeled with TMA (**Suppl. Figure 11**). In the PA-adapted method, the percentage of solvent B increased gradually, reaching 31% at 62 minutes and 41% at 70 minutes, before jumping to 72.2% at 71 minutes. In contrast, the TMA-adapted gradient increased solvent B at a slower initial rate, reaching 15.5% at 10 minutes and 25% at 45 minutes, before aligning with the PA gradient at 72.2% from 70 minutes onward. This slower increase in solvent strength in the early phase allows better retention and separation of the more hydrophobic peptides modified with TMA.

### DDA method

For data-dependent acquisition (DDA), MS1 spectra (from m/z 300 to 1300) were acquired at a resolution of 60,000, an AGC target of 1e6, and a maximum injection time (max IT) of 200 ms. MS/MS spectra were acquired at a resolution of 15,000, an AGC target value of 2e5, with a max IT of 100 ms, a loop count of 20, and a 1.5-m/z isolation window. Precursor ions were fragmented using higher-energy collisional dissociation (HCD) at 30% normalized collision energy. The first fixed mass was set to 80 m/z and dynamic exclusion time to 10 s.

### PRM method

Parallel Reaction Monitoring (PRM) was performed while targeting a resolution of 60 000, an AGC of 1.25x10^5^ and a maximum injection time of 110 ms and 190 ms. Synthetic peptides containing the Kacme modification and a heavy arginine residue (^13^C_6_-^15^N_4_) were bought from JPT to determine the retention time of peptides bearing the dual modification at each lysine of histones H3 and H4. These peptides were derivatized with TMA. PRM analyses were performed by providing a list of target ions to be monitored, including both endogenous light peptides and heavy synthetic peptide counterparts (**Suppl. Table 2**).

For assessing heavy acetyl incorporation, we targeted the mass of M+1 of each validated peptide and adjusted the isolation window to 2.5 m/z while targeting a resolution of 60 000, an AGC of 1.25x10^5^ and a maximum injection time of 170 ms.

## DATA ANALYSIS

### Peptide and PTM identification

For DDA data, the RAW MS files were converted into MGF files using the in-house developed tool MGFBoost, which leaves intact the relative intensities of fragments. Data were then interpreted by Mascot using an in-house histone database^36^. Validation of identifications and quantification was done in Proline^37^: a Mascot cut-off score of 25 was applied and peptidoforms were quantified using apex-intensities of their chromatographic peaks.

### Data Processing and Statistical Methods

Skyline was used to extract the area under the curve of the MS1-detected precursors as well as MS2 transitions. For acme and ac relative abundance analyses, we specified in Skyline a heavy isotope modification that corresponds to the mass difference between acetylation and acme, which is 14.0158 u. In this case, the light version on Skyline counts for the acetylated peptide while the heavy version counts for the acme-modified version. Finally, for heavy acetylation incorporation, we specified heavy ac isotope modification in which it adds 2.00671 u to the modification so it is possible to pick the transitions in the heavy section of fragments containing the heavy acetylation. Additionally, we specified acquisition method DIA although PRM was performed. Else, Skyline would only pick fragments from the targeted precursors, which in this case were masses of M+1 isotopes. Finally, we only considered the transitions carrying the modification in order to be able to estimate the heavy acetylation carried by this transition depending on the heavy transition signal observed.

### Processing of Proline and Skyline outputs

Histograms were obtained for the relative abundance of TMA- and PA-modified peptides by running code from library HistonePTM: https://github.com/HijaziHassan/histonePTM.

## SUPPORTING INFORMATION

A Supplementary Information file contains 12 additional figures and 2 tables.

## ACKNOWLEDGMENTS

We thank our colleagues at EDyP team for excellent technical support in proteomic analyses. The proteomic experiments were supported by Agence Nationale de la Recherche (ANR) under projects ProFI (Proteomics French Infrastructure, ANR-10-INBS-08) and GRAL, a program from the Chemistry Biology Health (CBH) Graduate School of University Grenoble Alpes (ANR-17-EURE-0003). PhD funding to HH was from GRAL and to HA from Association Huntington France. This project was also funded by ANR-21-CE44-0035-01 and ANR-23-CE12-0036 to DP.

