## Supplementary figures and tables for "Investigating Histone Lysine Acetylation and Acetyl-methylation in Mouse Brain using Enhanced Trimethylacetic Anhydride Derivatization Protocol"

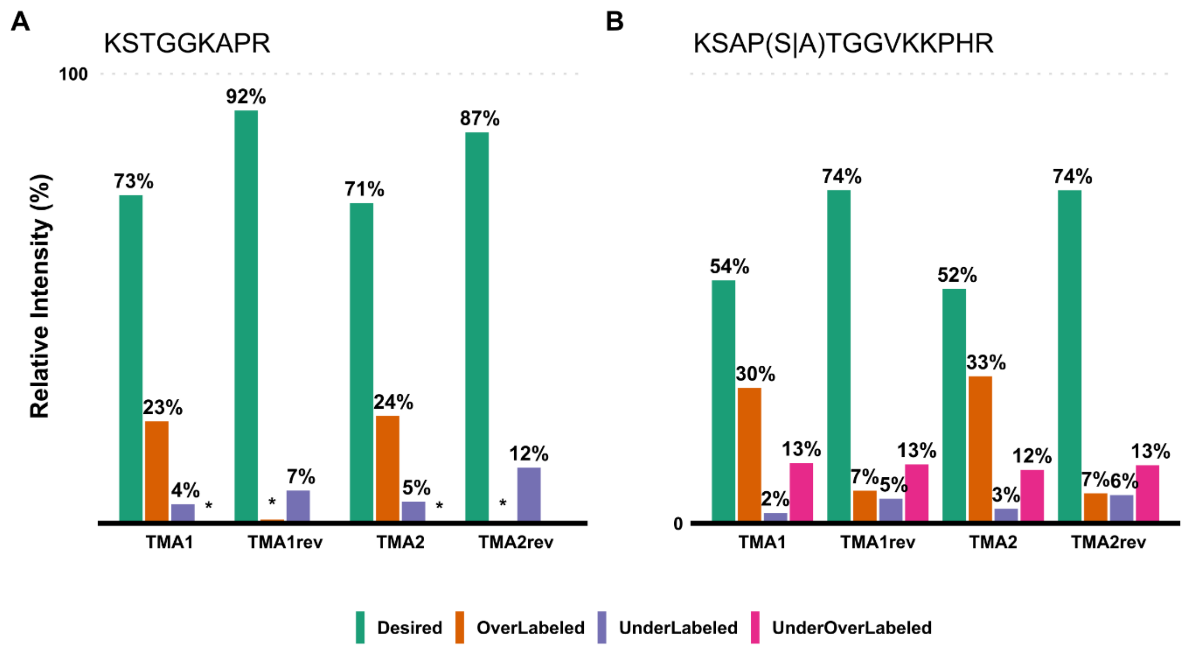


**Supplementary Figure 1. Estimation of the occurrence of over-TMAylation on Ser/Thr residues and of the efficiency of reversing this side reaction using a final step of basic treatment with hydroxylamine.** OverTMAylation assessment before (TMA1 and TMA2) and after one-hour hydroxylamine (HA; pH 12 at 500 mM) treatment (TMA1rev and TMArev2). The bars represent the summed relative abundance of H3 (9-17) peptidoforms H3 and H3.3 (27-40) that are fully modified at lysine residues and N-termini (desired), those that are TMAylated at Ser/Thr (OverLabeled) or those with failed TMAylation at the N-terminal and/or free or monomethylated Lys residues (UnderLabeled) or both (UnderOverLabeled). The values are rounded to the nearest integers. The asterisk indicates very low relative intensity of 1% or less. Its absence indicates that no peptide belonging to the corresponding category was identified. The graphs were generated using `ptm_labelingAssessment()` function from the *histonePTM* R package (https://github.com/HijaziHassan/histonePTM).

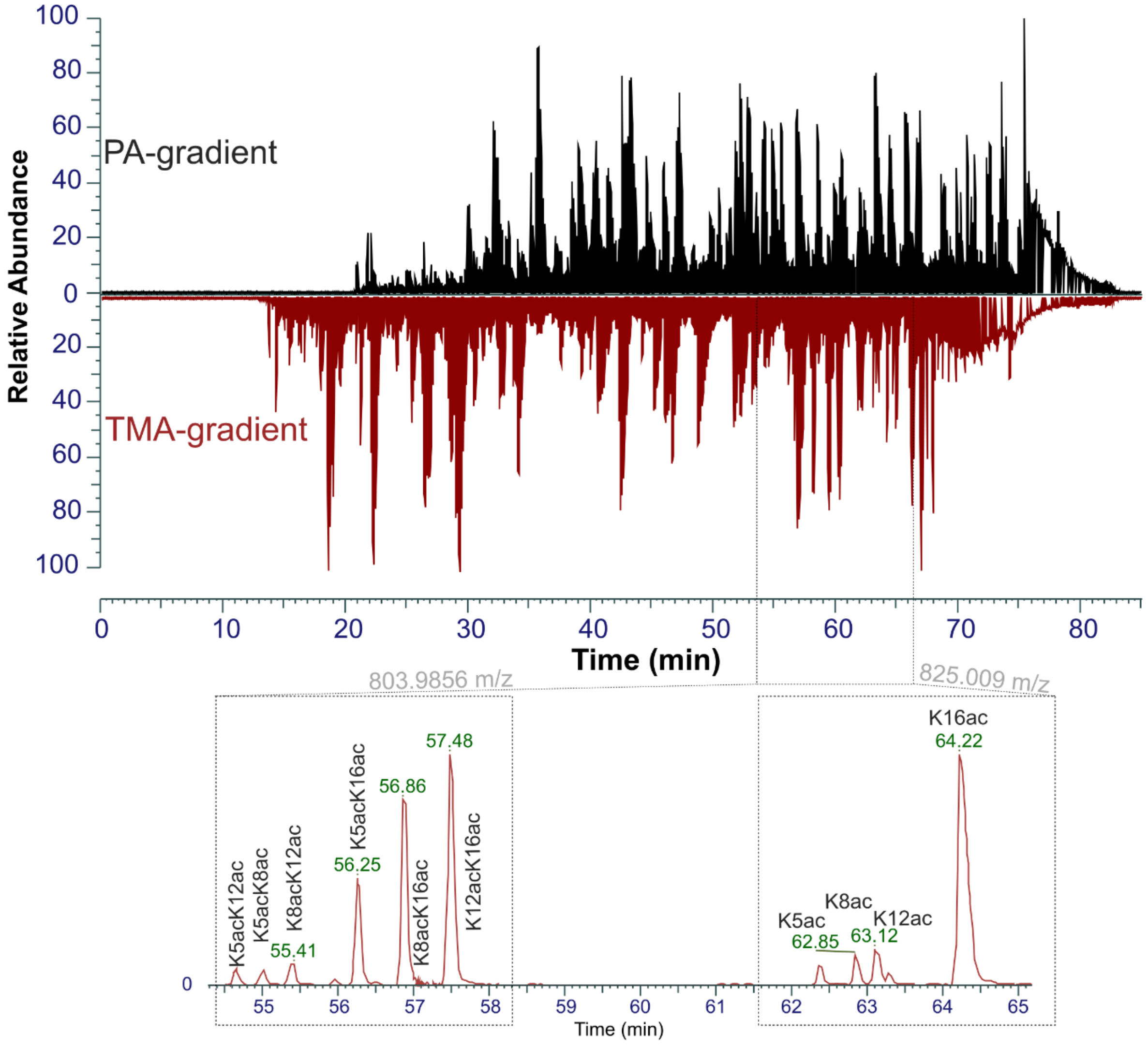


**Supplementary Figure 2. Impact of changing the elution gradient to adapt for increased hydrophobicity imparted by TMA compared to PA.** The distribution of TMAylated histone peptides over the whole elution time window was improved after modifying the gradient elution from the one used for propionylated histone peptides. The eXtracted Ion Chromatograms (XIC) of m/z 825.009 and 803.9856 corresponding to mono- and doubly-acetylated positional isomers of $G_4$KGGKGLGKGGAK$R_{17}$ peptide covering the N-terminal tail of H4, which contains the modifiable lysines K5, K8, K12 and K16, reveal their complete chromatographic separation. PA: propionic anhydride; TMA: trimethylacetic anhydride.

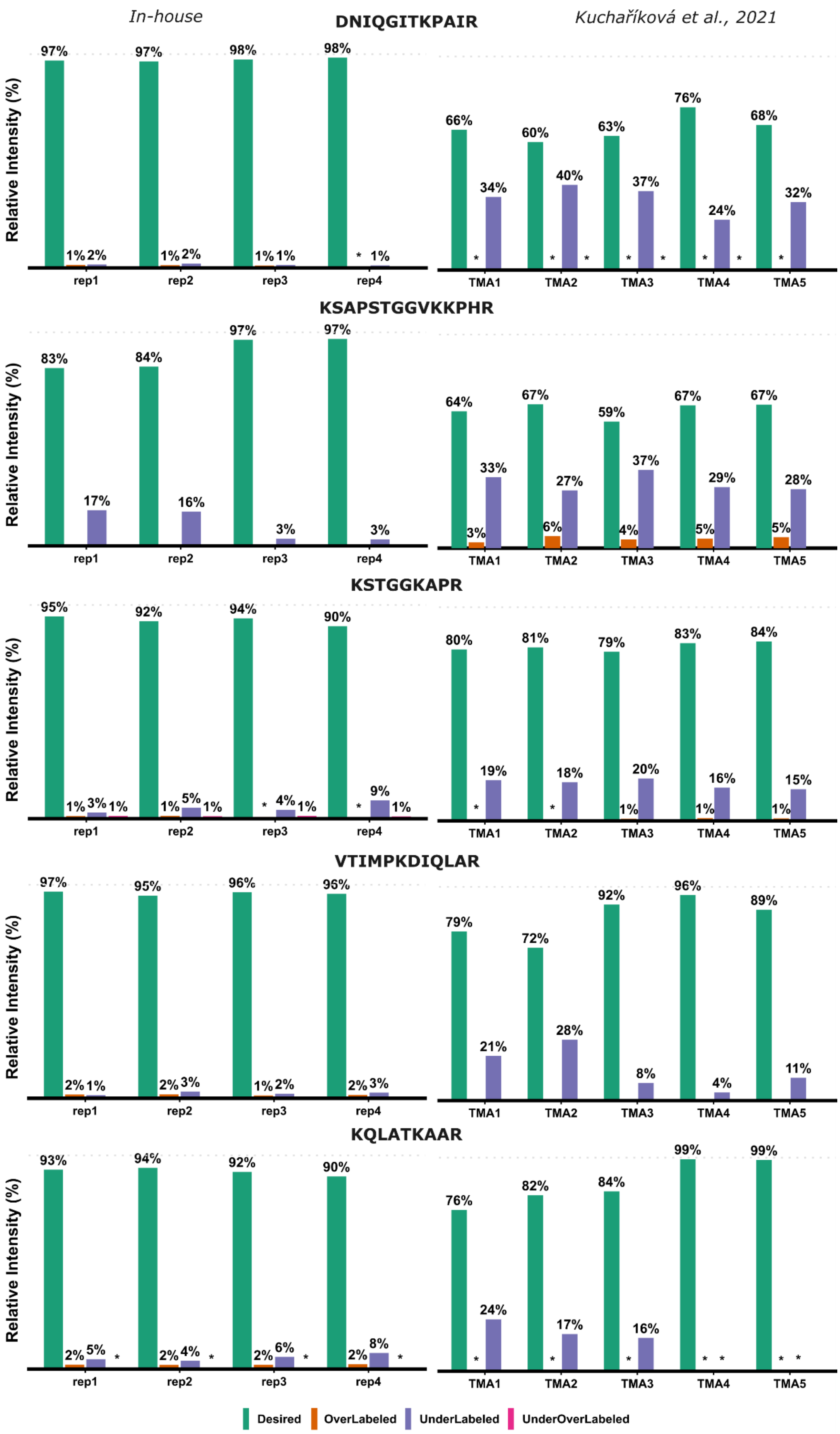
In-house
DNIQGITKPAIR
Kuchaříková et al., 2021
KSAPSTGGVKKPHR
KSTGGKAPR
VTIMPKDIQLAR
KQLATKAAR
Relative Intensity (%)
rep1
rep2
rep3
rep4
TMA1
TMA2
TMA3
TMA4
TMA5
Desired
OverLabeled
UnderLabeled
UnderOverLabeled

**Supplementary Figure 3. A comparison between our optimized TMAylation protocol and the originally published one (Kuchaříková et al.).** The raw files were searched with Mascot while specifying as variable modifications TMA, me1, me1-TMA, ac, me2, me3 on lysines, and TMA (any peptide N-terminal). Filtering with a minimal Mascot score and quantification were done in Proline. Raw files acquired on control samples derivatized with TMA (i.e. Lumos1_105min-IS_TMA_ctrl_1-5.raw) were downloaded from PRIDE (PXD019502) and were named TMA1 to TMA5. The asterisk indicates very low relative intensities of 1% or less. Its absence indicates that no peptide belonging to the corresponding category was identified. The graphs were generated using `ptm_labelingAssessment()` function from the *histonePTM* R package (https://github.com/HijaziHassan/histonePTM).

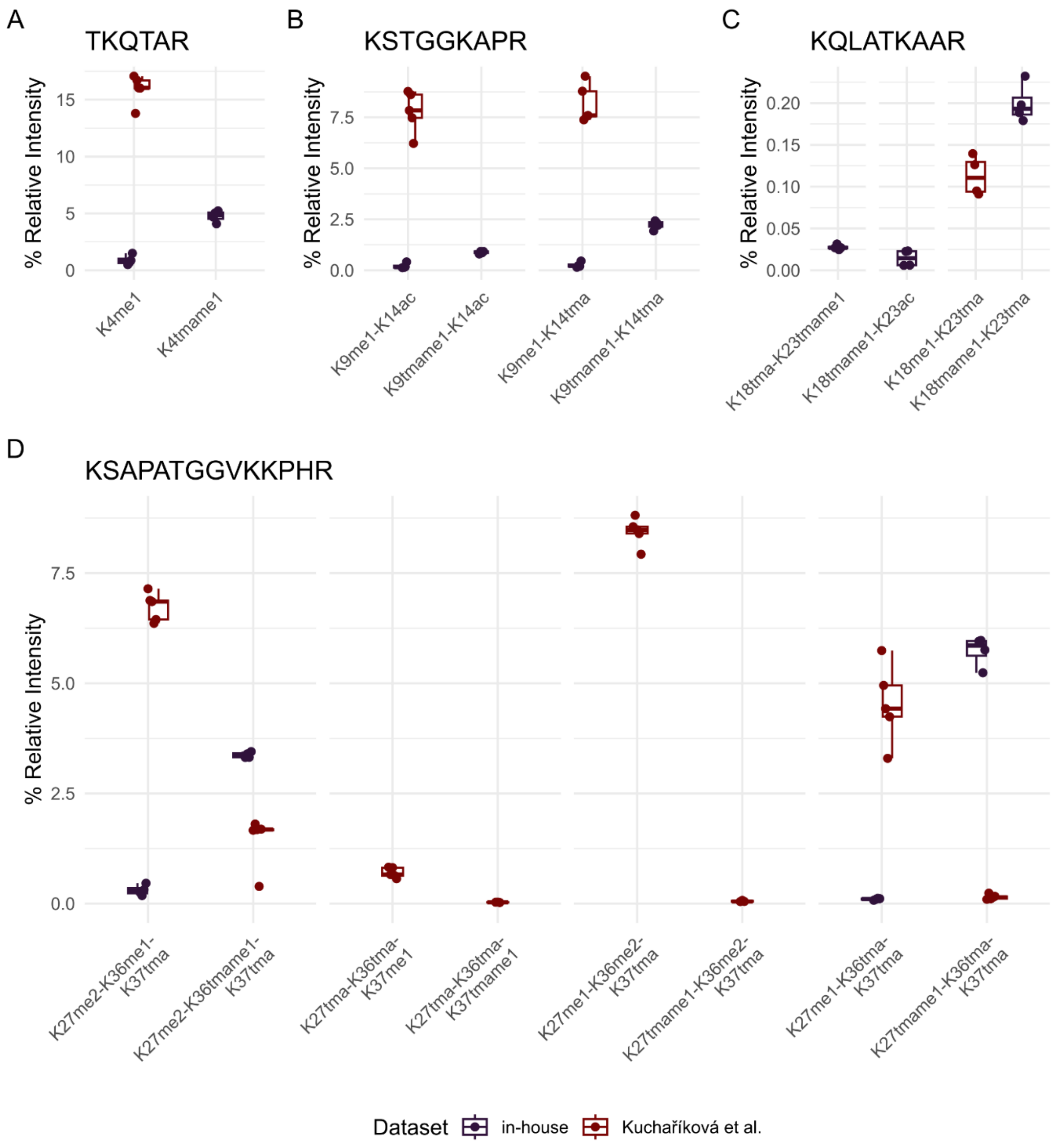


**Supplementary Figure 4. Comparison of labeling efficiency of monomethylated lysines between our improved TMAylation protocol and the original protocol (Kuchaříková et al.).** The box plots with jittered points represent the relative MS intensities of H3 N-terminal peptides that failed to be labeled (me1) or were successfully labeled (tmame1) (n=4). The aim was not to compare intensities inter-samples but rather intra-samples as histones were extracted from different cell types. The comparison was done over peptides that were successfully N-terminally TMAylated. The graphs were generated using `quant_me1LabelingPlot()` function from the *histonePTM* R package (https://github.com/HijaziHassan/histonePTM).

**A**

**H2A.X**

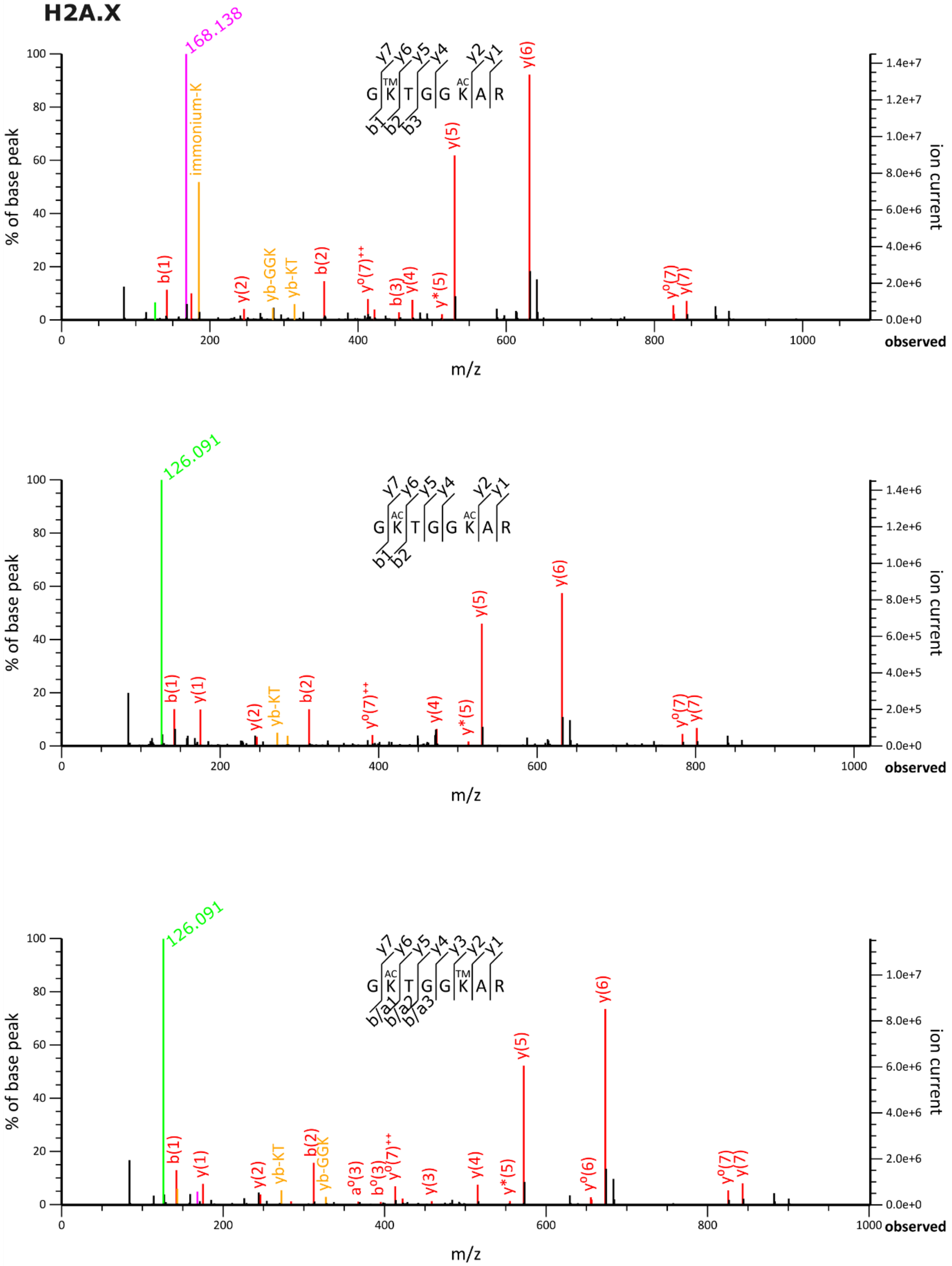
168.138
immonium-K
y7 y6 y5 y4 y2 y1
G K T G G K A R
TM
AC
b1 b2 b3
y(6)
y(5)
b(1)
y(2)
yb-GGK
yb-KT
b(2)
y°(7)++
b(3)
y(4)
y*(5)
y°(7)
y(7)
% of base peak
ion current
m/z
observed
126.091
y7 y6 y5 y4 y2 y1
G K T G G K A R
AC
AC
b1 b2
y(6)
y(5)
b(1)
y(1)
y(2)
yb-KT
b(2)
y°(7)++
y(4)
y*(5)
y°(7)
y(7)
% of base peak
ion current
m/z
observed
126.091
y7 y6 y5 y4 y3 y2 y1
G K T G G K A R
AC
TM
b/a1 b/a2 b/a3
y(6)
y(5)
b(1)
y(1)
y(2)
yb-KT
b(2)
yb-GGK
a°(3)
b°(3)
y°(7)++
y(3)
y(4)
y*(5)
y°(6)
y°(7)
y(7)
% of base peak
ion current
m/z
observed

**B**

**H2A**

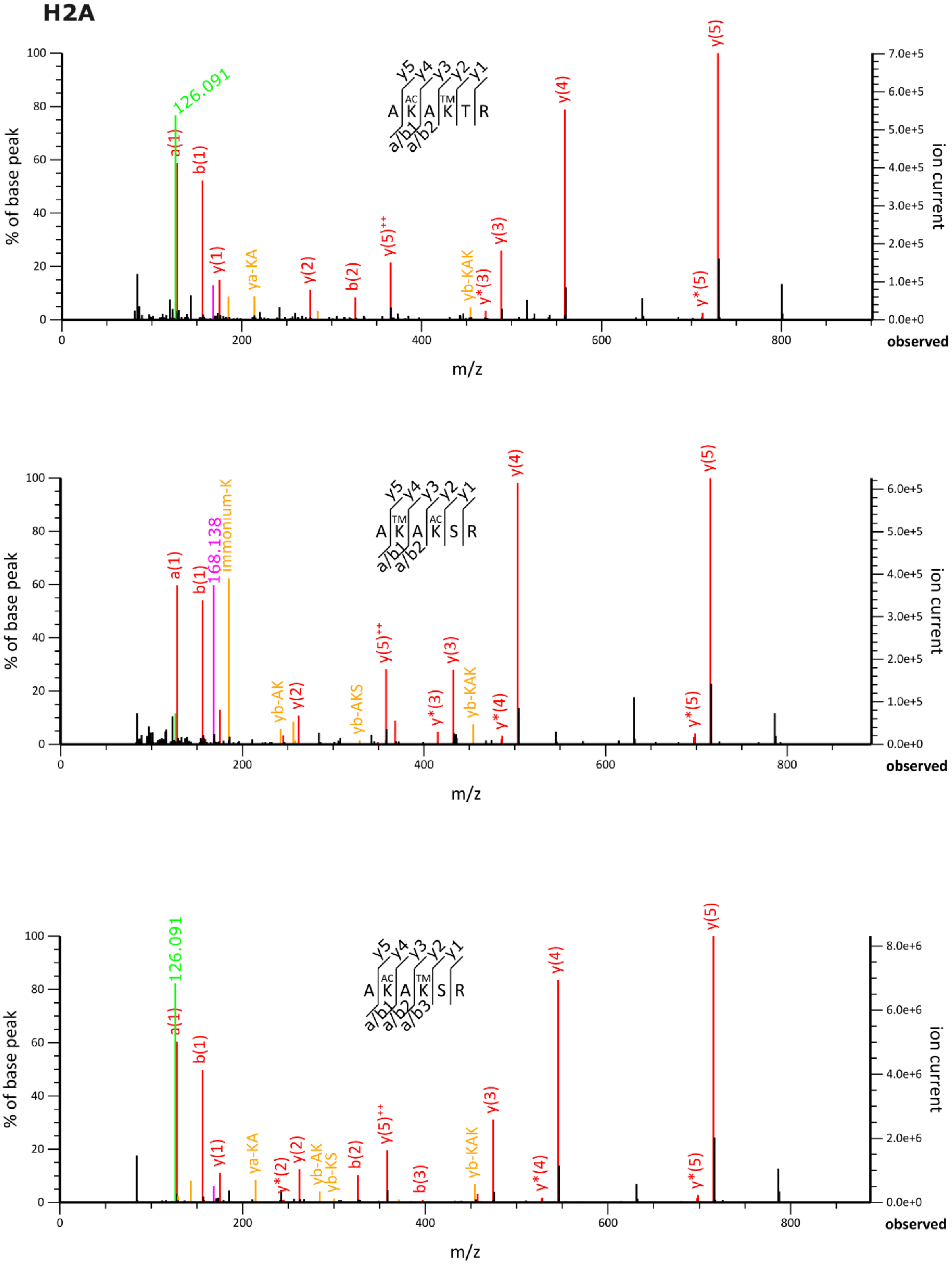

A K(AC) A K(TM) T R
y5 y4 y3 y2 y1
a/b1 a/b2
126.091
a(1)
b(1)
y(1)
ya-KA
y(2)
b(2)
y(5)++
yb-KAK
y*(3)
y(3)
y(4)
y*(5)
y(5)
A K(TM) A K(AC) S R
168.138
Immonium-K
yb-AK
yb-AKS
y*(4)
A K(AC) A K(TM) S R
a/b3
y*(2)
yb-KS
b(3)
% of base peak
ion current
m/z
observed

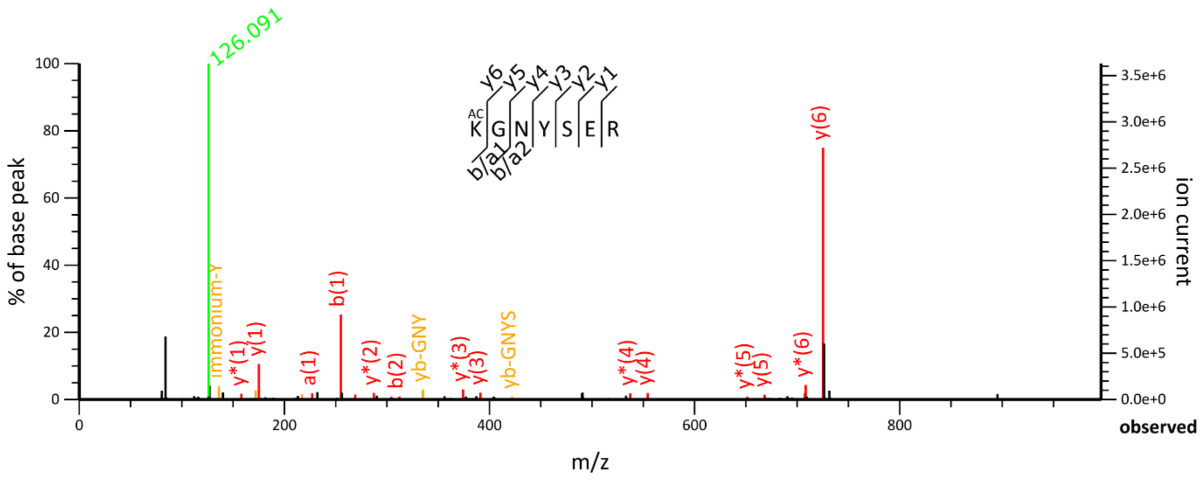
126.091
KGNYSER
y6 y5 y4 y3 y2 y1
b/a1 b/a2
AC
y(6)
b(1)
immonium-Y
y*(1)
y(1)
a(1)
y*(2)
b(2)
yb-GNY
y*(3)
y(3)
yb-GNYS
y*(4)
y(4)
y*(5)
y(5)
y*(6)
% of base peak
ion current
m/z
observed

**C**
**H2B**

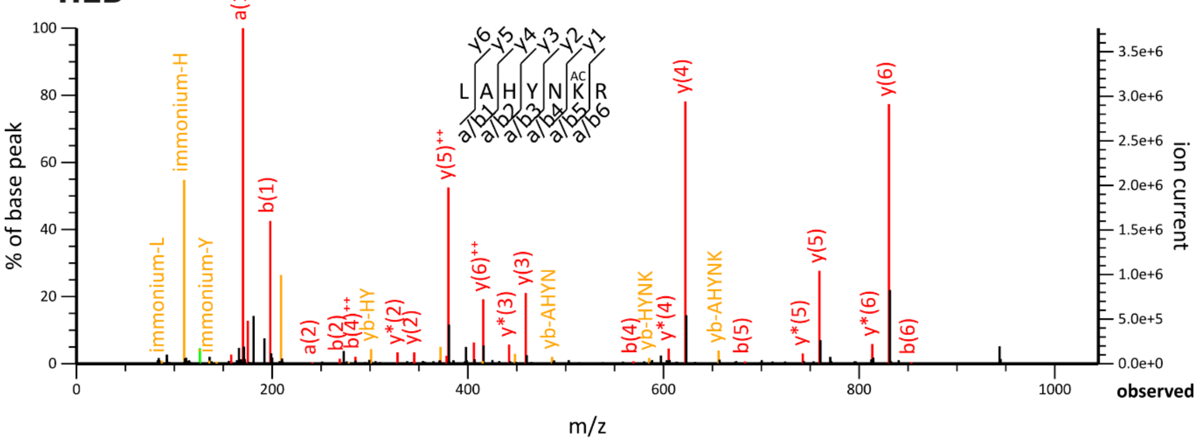
LAHYNKR
y6 y5 y4 y3 y2 y1
a/b1 a/b2 a/b3 a/b4 a/b5 a/b6
AC
a(1)
immonium-H
immonium-L
immonium-Y
b(1)
y(5)++
y(6)++
y(3)
y(4)
y(6)
y(5)
a(2)
b(2)
b(4)++
yb-HY
y*(2)
y(2)
y*(3)
yb-AHYN
b(4)
yb-HYNK
y*(4)
yb-AHYNK
b(5)
y*(5)
y*(6)
b(6)
% of base peak
ion current
m/z
observed

**D**
**H1**

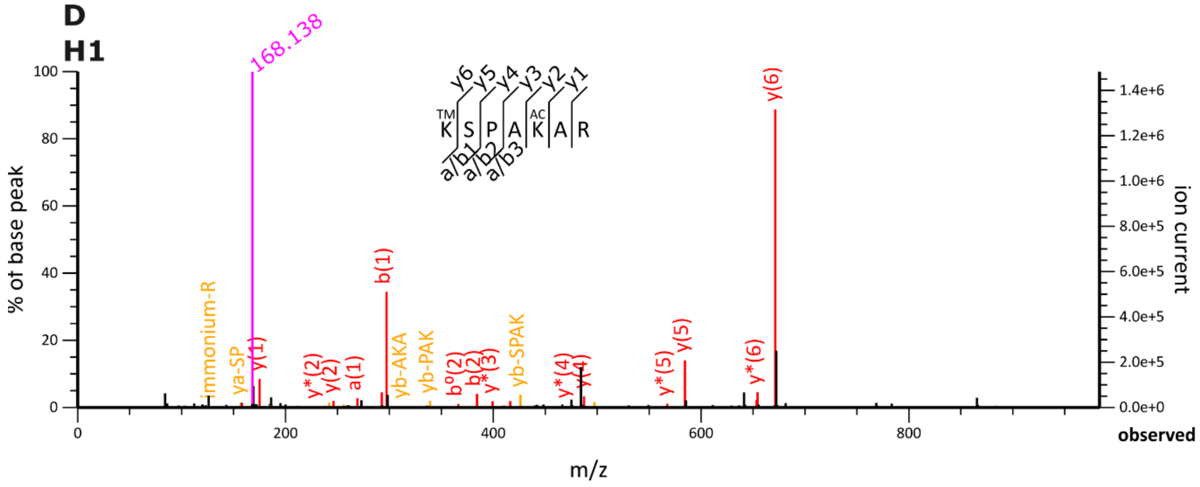
168.138
KSPAKAR
y6 y5 y4 y3 y2 y1
a/b1 a/b2 a/b3
TM
AC
y(6)
b(1)
immonium-R
ya-SP
y(1)
y*(2)
y(2)
a(1)
yb-AKA
yb-PAK
b°(2)
b(2)
y*(3)
yb-SPAK
y*(4)
y(4)
y*(5)
y(5)
y*(6)
% of base peak
ion current
m/z
observed

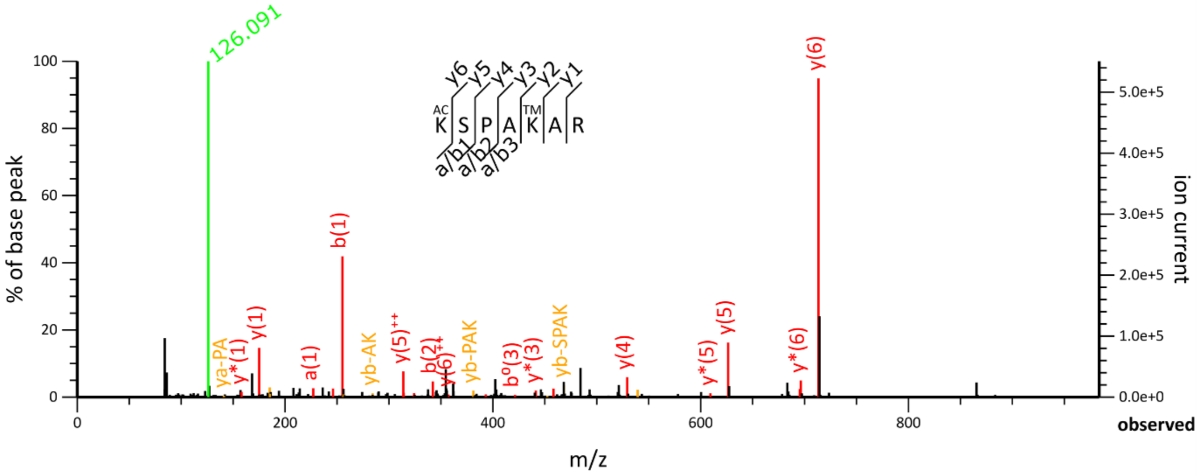

126.091
y(6)
b(1)
y(5)
y(1)
y*(1)
ya-PA
a(1)
yb-AK
y(5)++
b(2)
y(6)++
yb-PAK
b°(3)
y*(3)
yb-SPAK
y(4)
y*(5)
y*(6)
% of base peak
ion current
m/z
observed


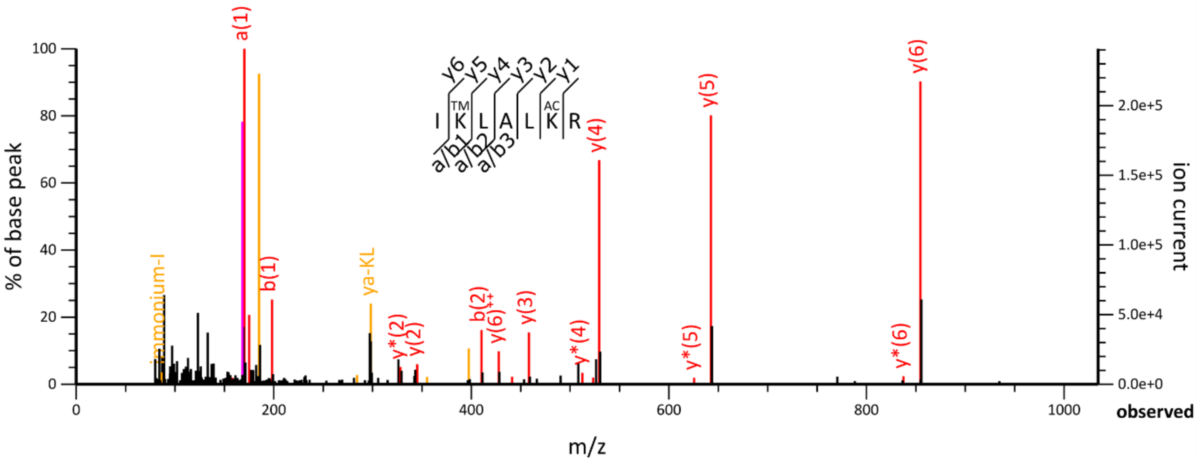

a(1)
y(6)
y(5)
y(4)
b(1)
Immonium-I
ya-KL
y*(2)
y(2)
b(2)
y(6)++
y(3)
y*(4)
y*(5)
y*(6)
% of base peak
ion current
m/z
observed


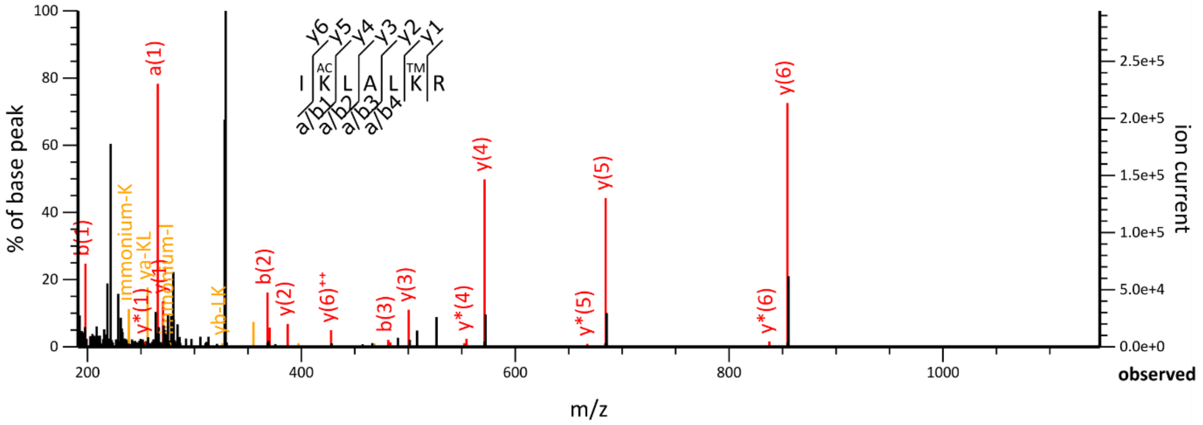

a(1)
y(6)
y(4)
y(5)
b(1)
Immonium-K
ya-KL
y*(1)
yb-LK
b(2)
y(2)
y(6)++
b(3)
y(3)
y*(4)
y*(5)
y*(6)
% of base peak
ion current
m/z
observed

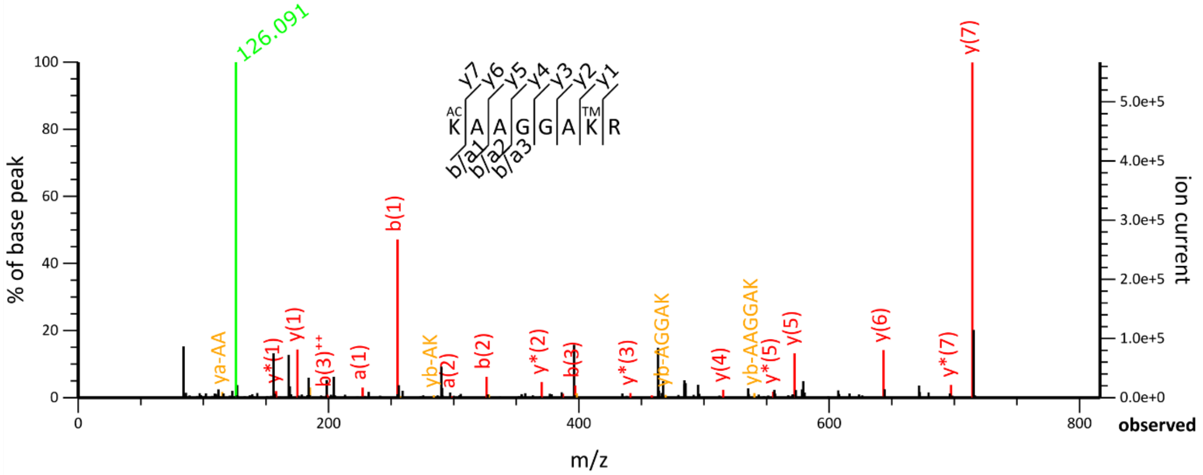


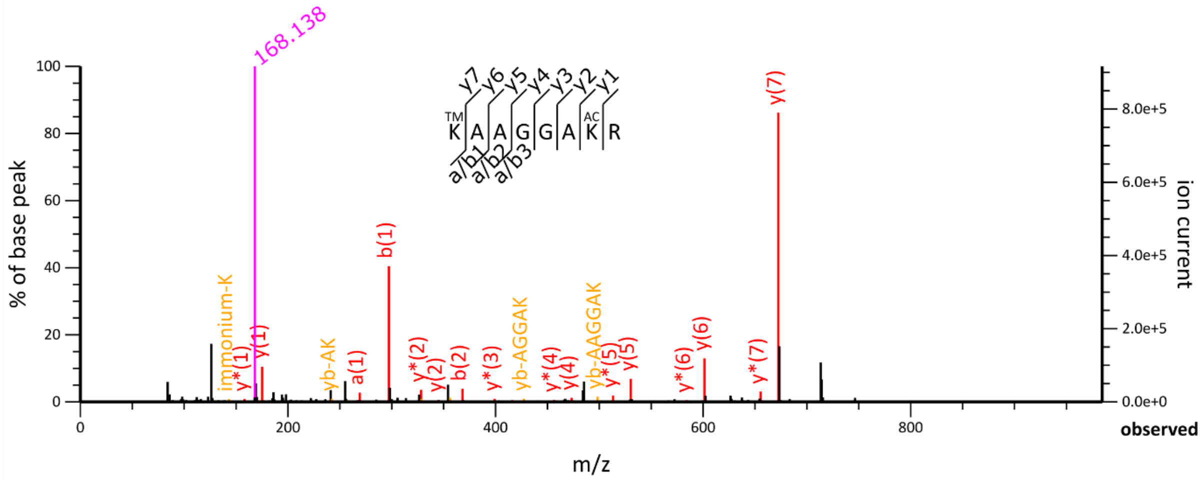


**E**

### H1.6 mouse-specific

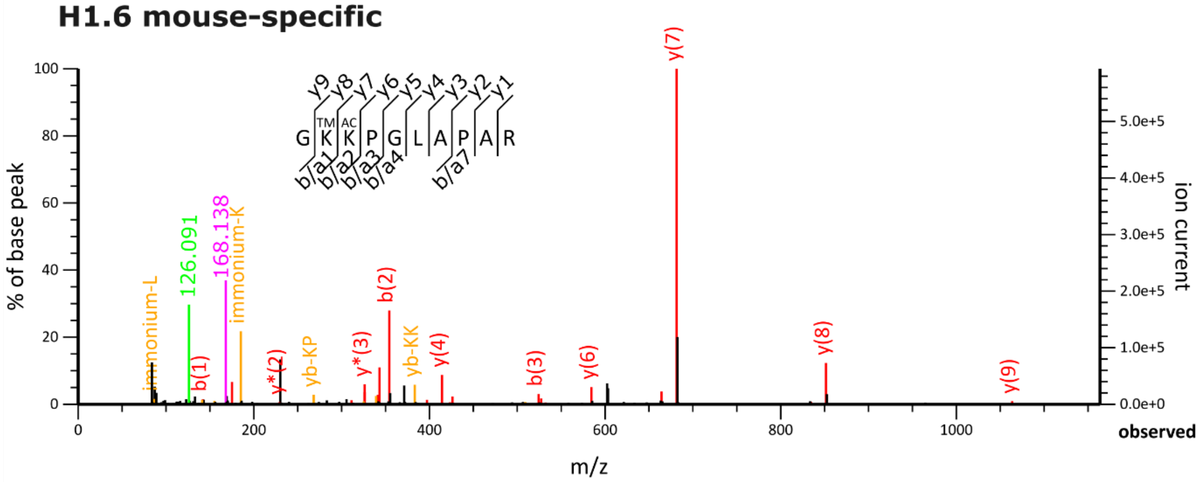

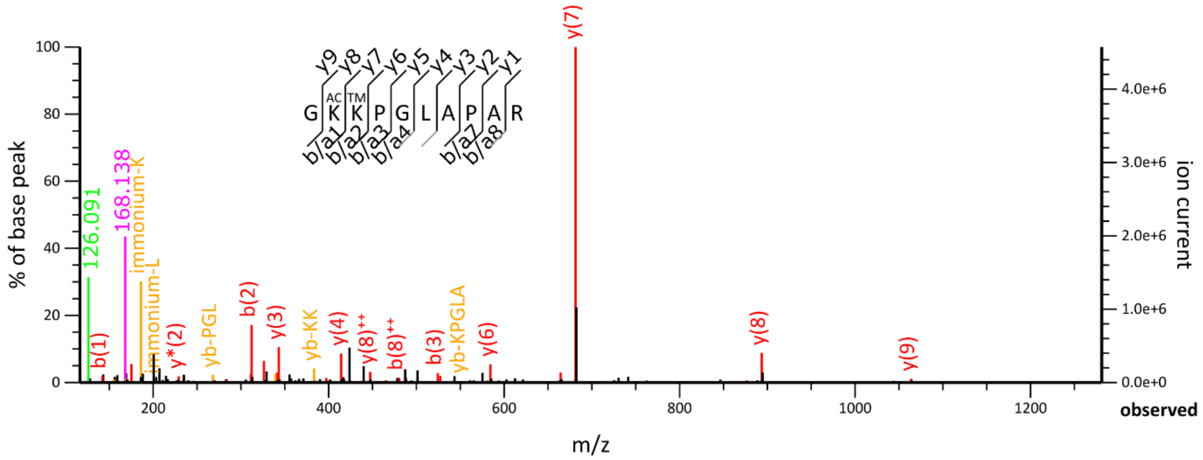

G K K P G L A P A R
AC TM
y9 y8 y7 y6 y5 y4 y3 y2 y1
b/a1 b/a2 b/a3 b/a4 b/a7 b/a8
126.091
168.138
immonium-K
immonium-L
b(1)
y*(2)
yb-PGL
b(2)
y(3)
yb-KK
y(4)
y(8)++
b(8)++
b(3)
yb-KPGLA
y(6)
y(7)
y(8)
y(9)
% of base peak
ion current
m/z
observed


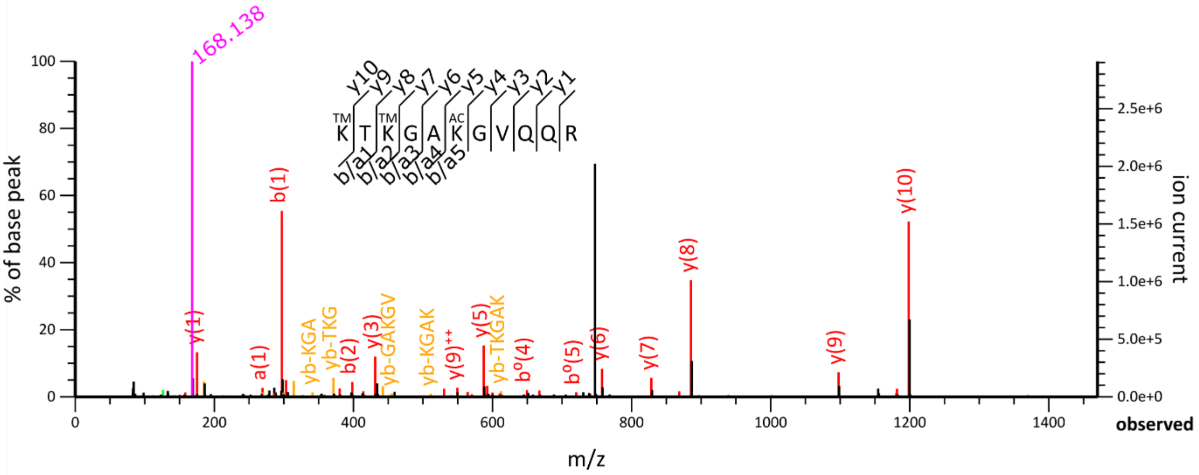

K T K G A K G V Q Q R
TM TM AC
y10 y9 y8 y7 y6 y5 y4 y3 y2 y1
b/a1 b/a2 b/a3 b/a4 b/a5
168.138
b(1)
y(1)
a(1)
yb-KGA
yb-TKG
y(3)
b(2)
yb-GAKGV
yb-KGAK
y(9)++
y(5)
yb-TKGAK
b°(4)
b°(5)
y(6)
y(7)
y(8)
y(9)
y(10)
% of base peak
ion current
m/z
observed


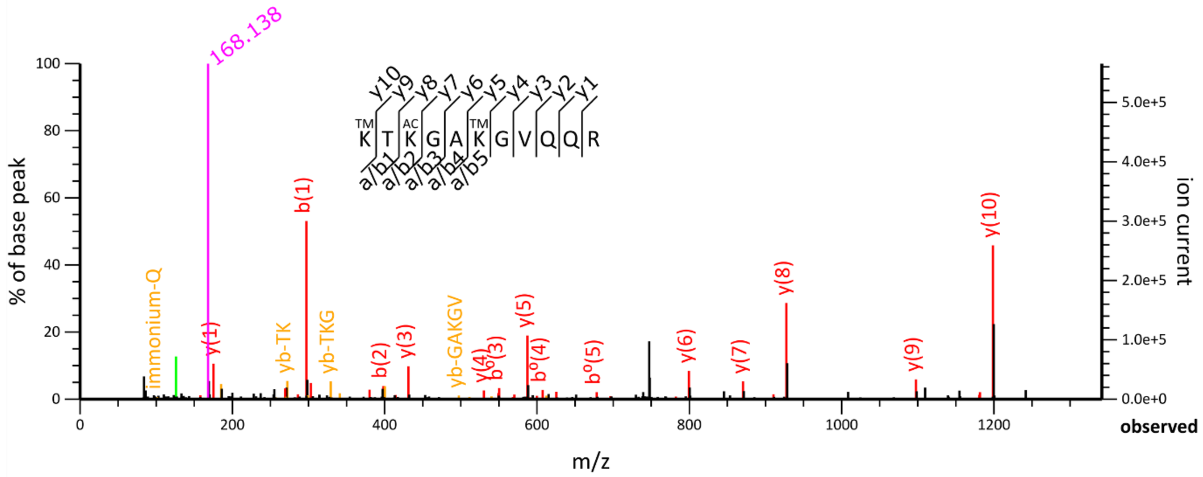

K T K G A K G V Q Q R
TM AC TM
y10 y9 y8 y7 y6 y5 y4 y3 y2 y1
a/b1 a/b2 a/b3 a/b4 a/b5
168.138
immonium-Q
y(1)
yb-TK
b(1)
yb-TKG
b(2)
y(3)
yb-GAKGV
y(4)
b°(3)
y(5)
b°(4)
b°(5)
y(6)
y(7)
y(8)
y(9)
y(10)
% of base peak
ion current
m/z
observed

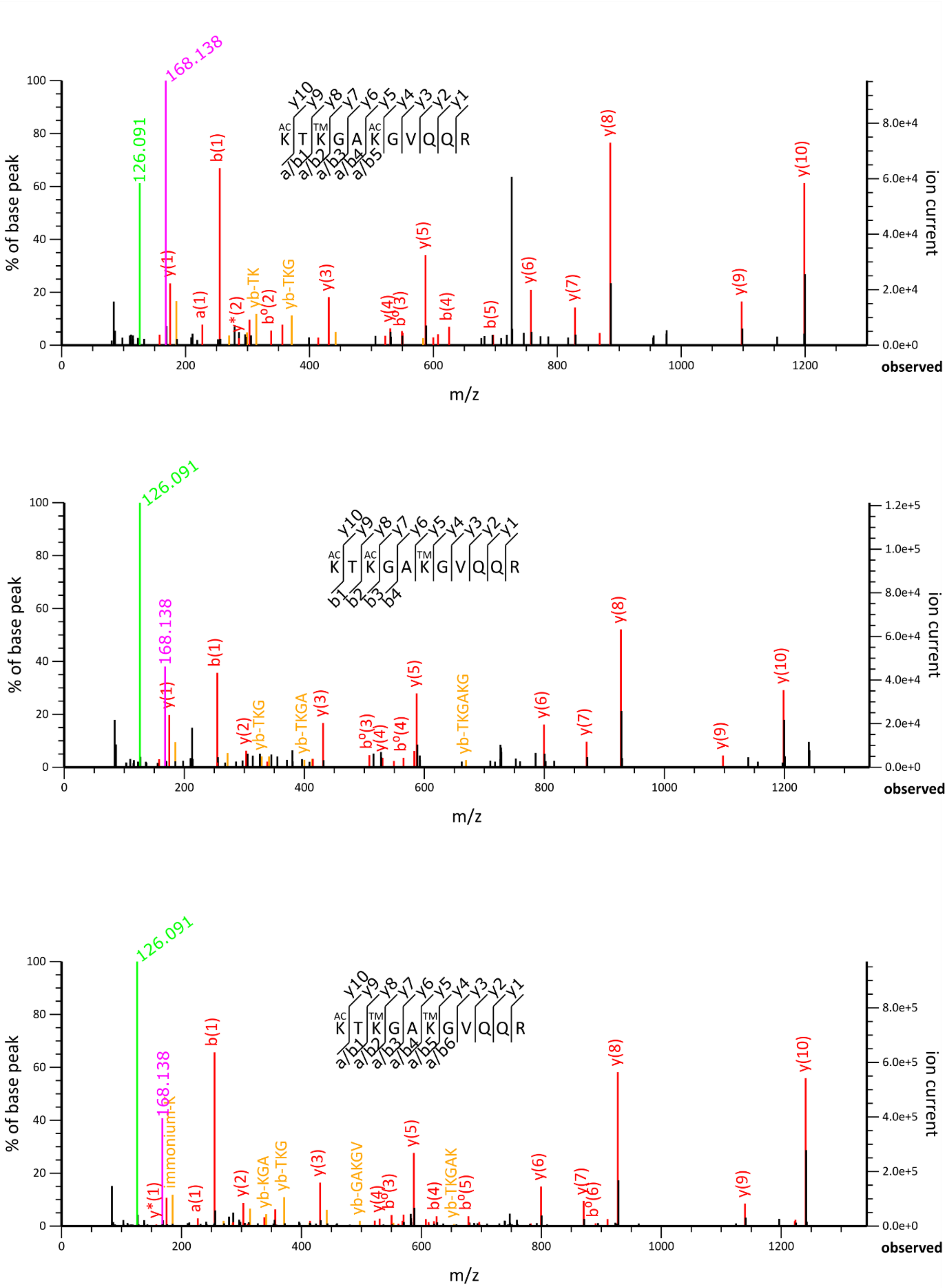

100
80
60
40
20
0
% of base peak
ion current
8.0e+4
6.0e+4
4.0e+4
2.0e+4
0.0e+0
0
200
400
600
800
1000
1200
observed
m/z
168.138
126.091
b(1)
y(1)
a(1)
y*(2)
yb-TK
b°(2)
yb-TKG
y(3)
y(4)
b°(3)
y(5)
b(4)
b(5)
y(6)
y(7)
y(8)
y(9)
y(10)
1.2e+5
1.0e+5
8.0e+4
6.0e+4
4.0e+4
2.0e+4
0.0e+0
y(2)
yb-TKGA
b°(3)
b°(4)
yb-TKGAKG
8.0e+5
6.0e+5
4.0e+5
2.0e+5
0.0e+0
immonium-K
y*(1)
yb-KGA
yb-GAKGV
yb-TKGAK
b°(5)
b°(6)

**F**

**H3.3**

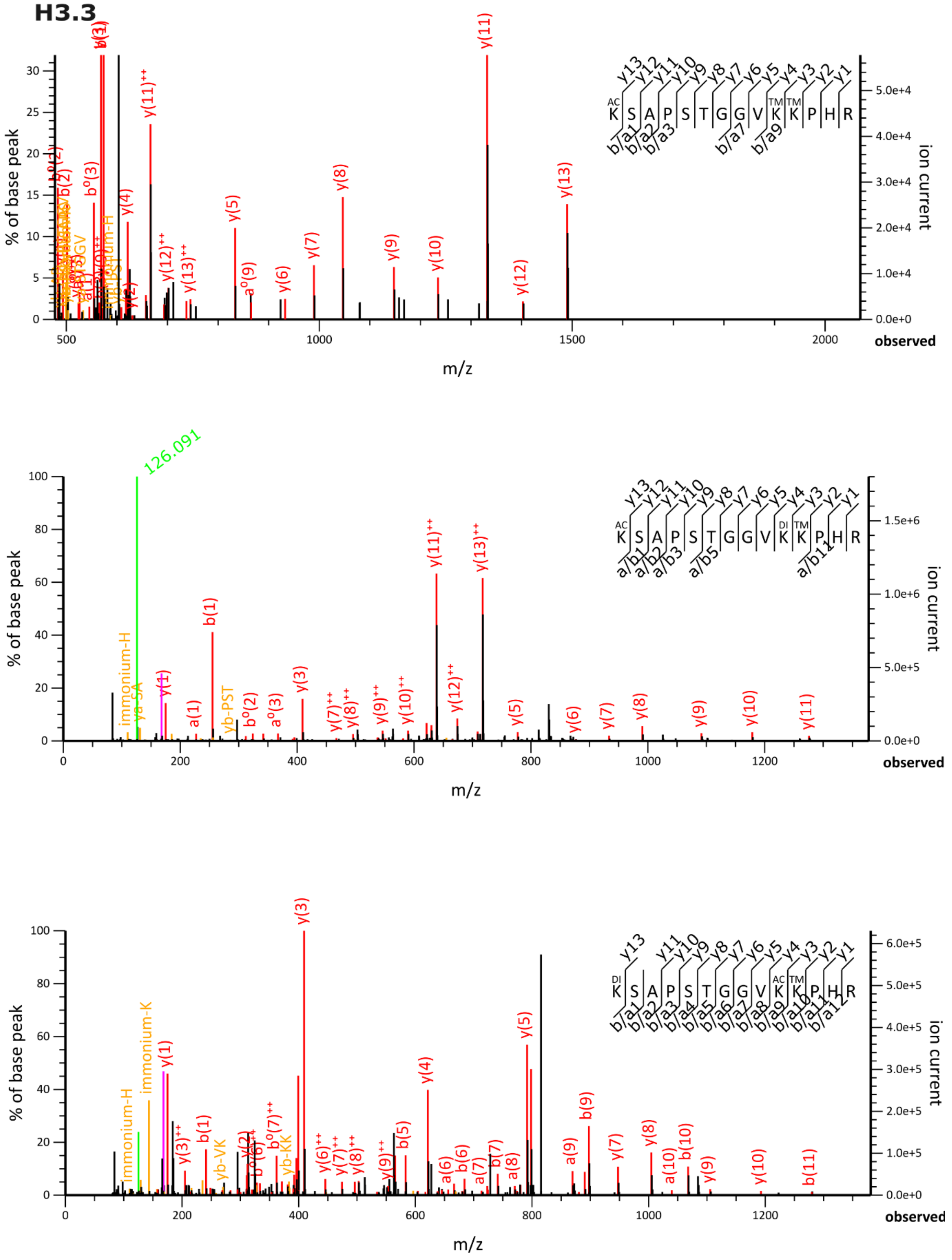

% of base peak
ion current
m/z
observed
K S A P S T G G V K K P H R
y(11)
y(13)
y(8)
y(11)++
y(13)++
126.091
b(1)
y(3)
y(5)
immonium-H
immonium-K
y(1)
y(4)
b(9)

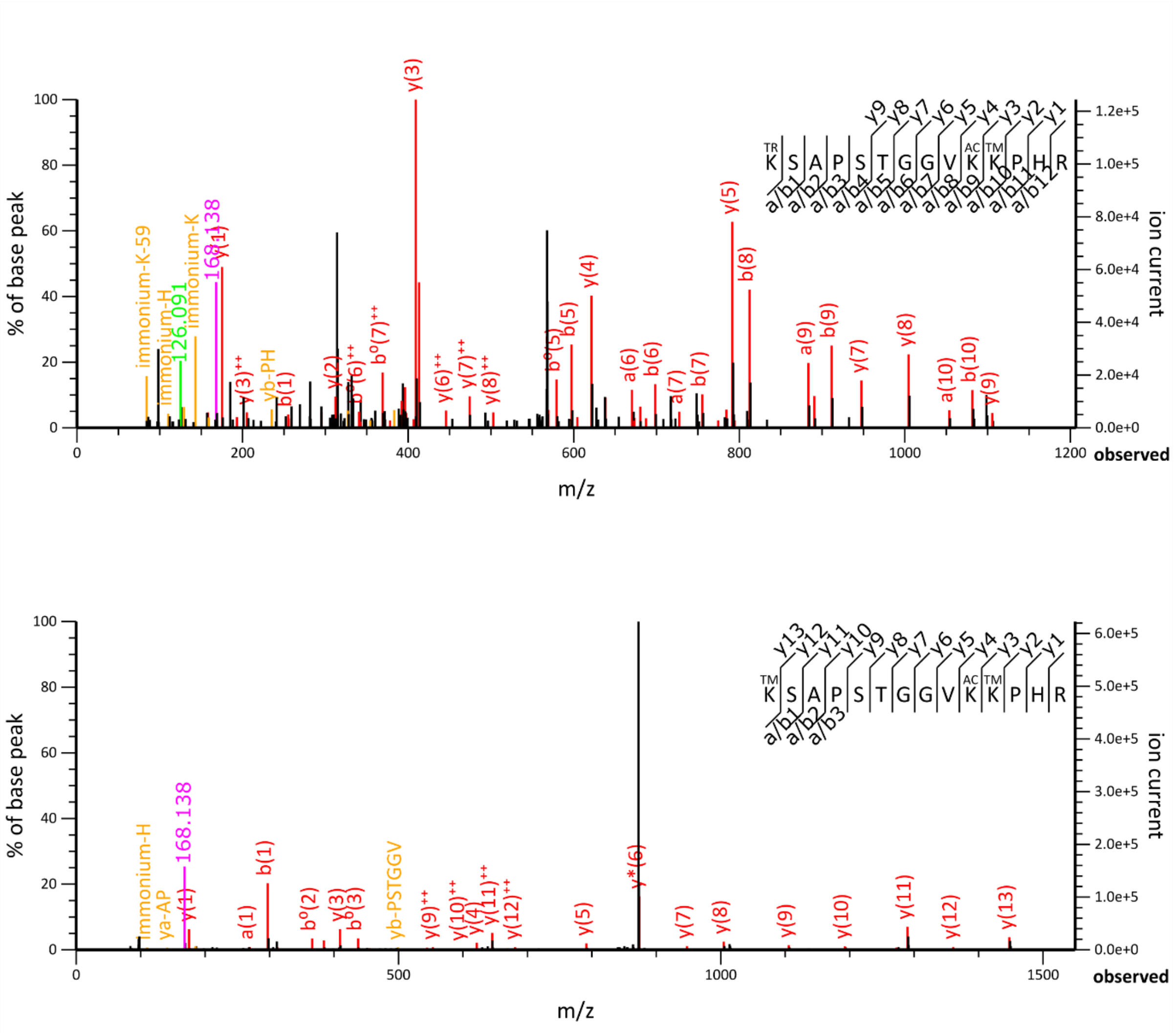


**Supplementary Figure 5. TMA-derivatization of histone peptides enhances the detection of acetylated peptides.** Several MS/MS spectra representing acetylated peptides identified in samples treated with trimethylacetic anhydride from histone proteins: **(A)** H2A, **(B)** H2A.X, **(C)** H2B, **(D)** H1, **(E)** mouse-specific H1.6, and **(F) H3.3**. Trimethylacetic anhydride (N-terminal; +84.0575 Da) was set as static modification. TM: Trimethylacetylation; AC: Acetylation; TR: Trimethylation; DI: Dimethylation. Cyclic immonium (CycIm) ions of Kac (126.091 m/z) and Ktm (168.138 m/z) are colored in bright green and fuchsia, respectively.

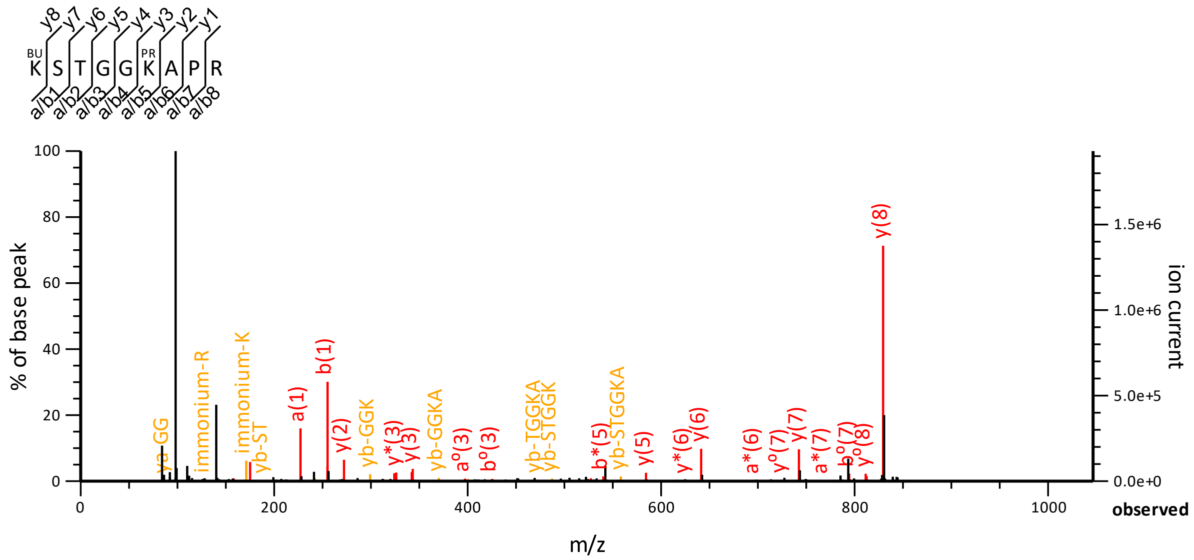

K S T G G K A P R
% of base peak
ion current
m/z
observed
immonium-R
immonium-K
yb-ST
a(1)
b(1)
y(2)
yb-GGK
yb-GGKA
y(3)
yb-TGGKA
yb-STGGK
yb-STGGKA
y(5)
y(6)
y(7)
y(8)


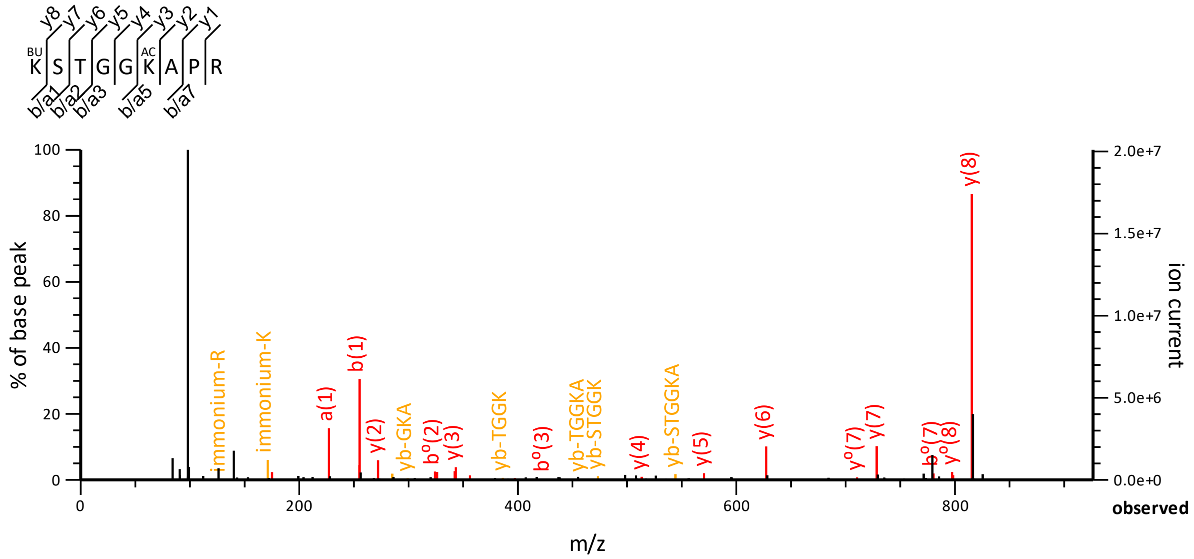

K S T G G K A P R
% of base peak
ion current
m/z
observed
immonium-R
immonium-K
a(1)
b(1)
y(2)
yb-GKA
y(3)
yb-TGGK
yb-TGGKA
yb-STGGK
y(4)
yb-STGGKA
y(5)
y(6)
y(7)
y(8)

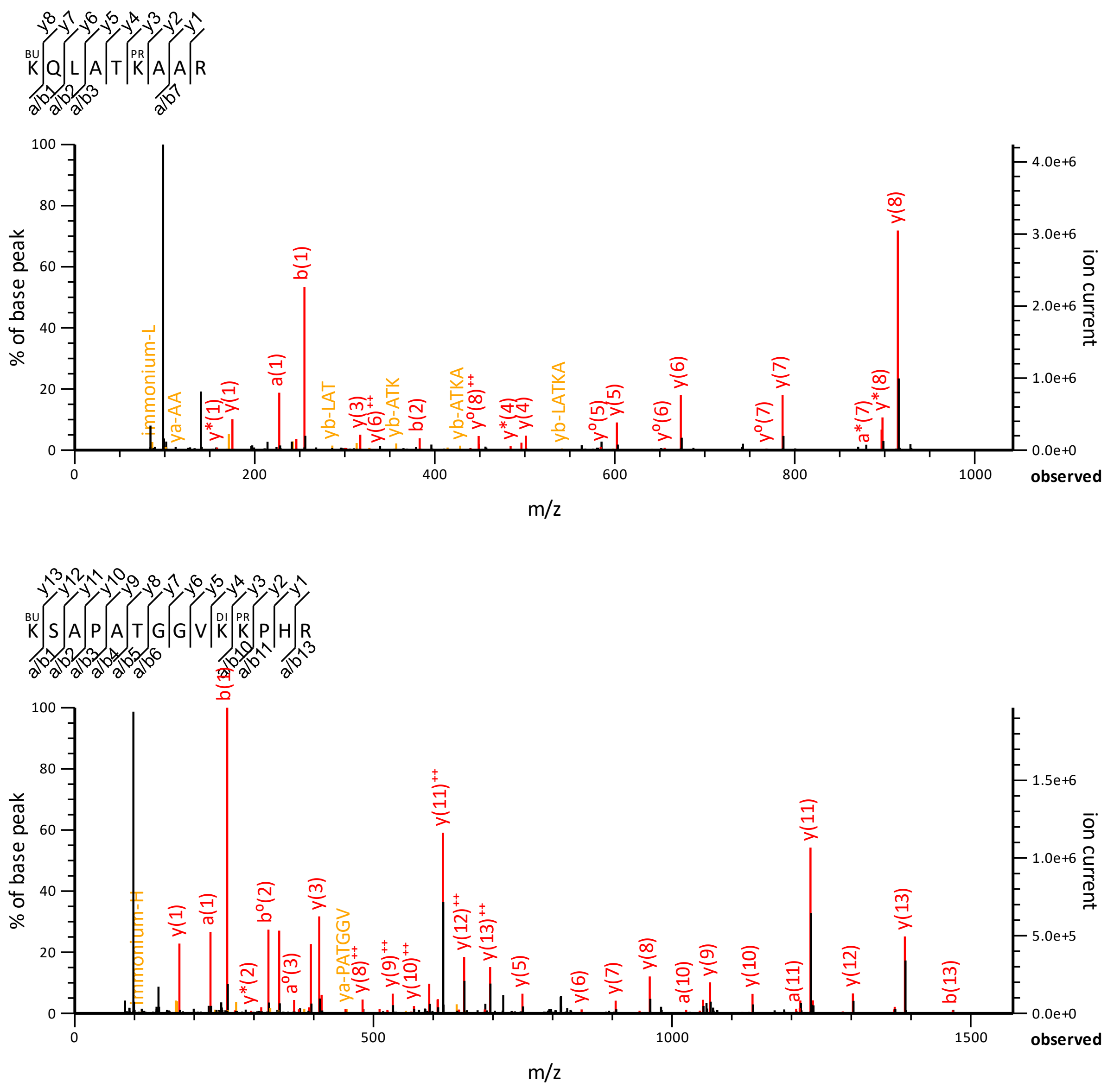

**Supplementary Figure 6: MS/MS spectra identifying peptides that contain a Kme1pr residue in first position (labeled as Kbu by the program Mascot), and exhibiting an intense CycIm at m/z 98.097 characteristic of Kme1.**

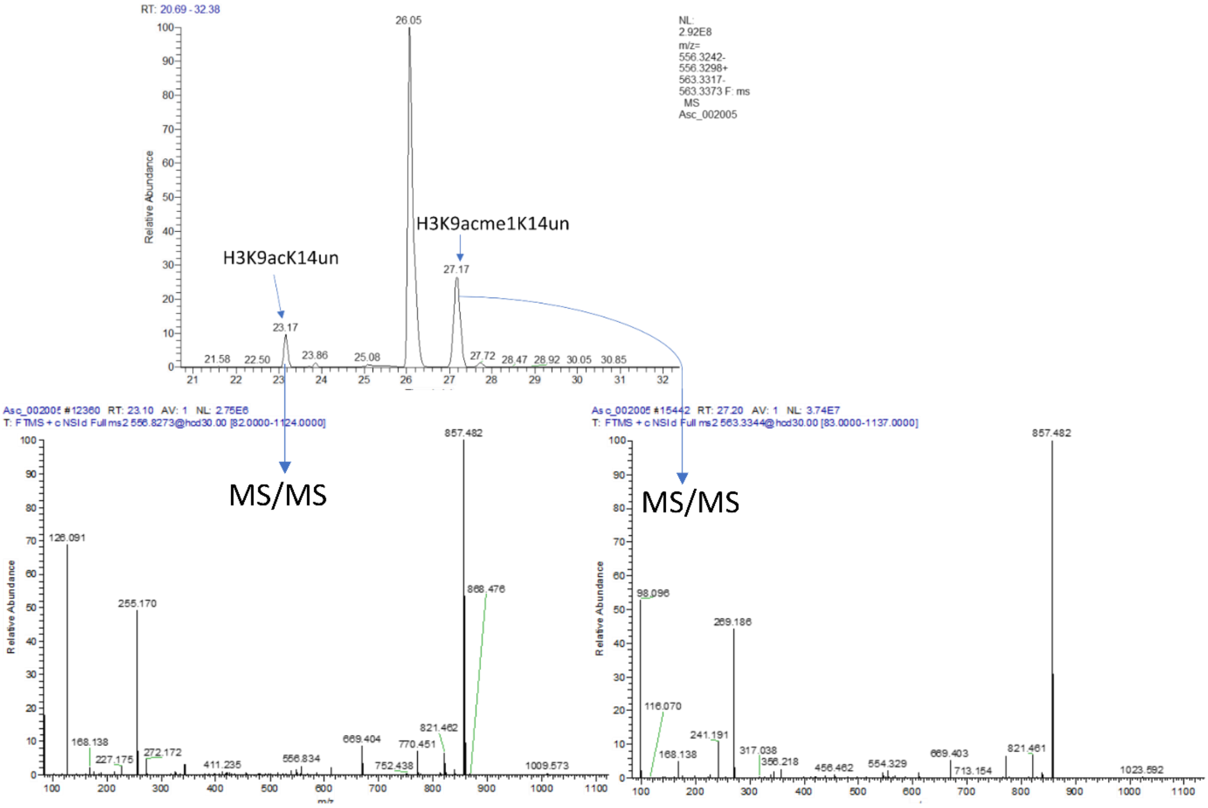


**Supplementary Figure 7: eXtracted Ion Chromatograms (XIC) of m/z 556.327 and 563.3345 corresponding to peptide H3 K9STGGK14APR acetylated or modified by acme at its first position.** These relative signals were observed in first DDA analyses of histones extracted from mouse brain and modified by TMA.

| Valeric Anhydride | Trimethylacetic Anhydride |
|---|---|
| 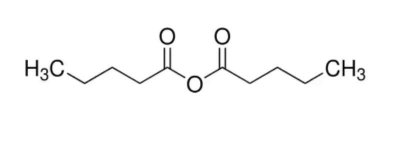 | 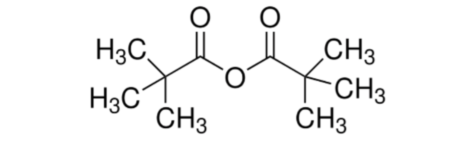 |

**Supplementary Figure 8: Structures of valeric anhydride and trimethylacetic anhydride**.

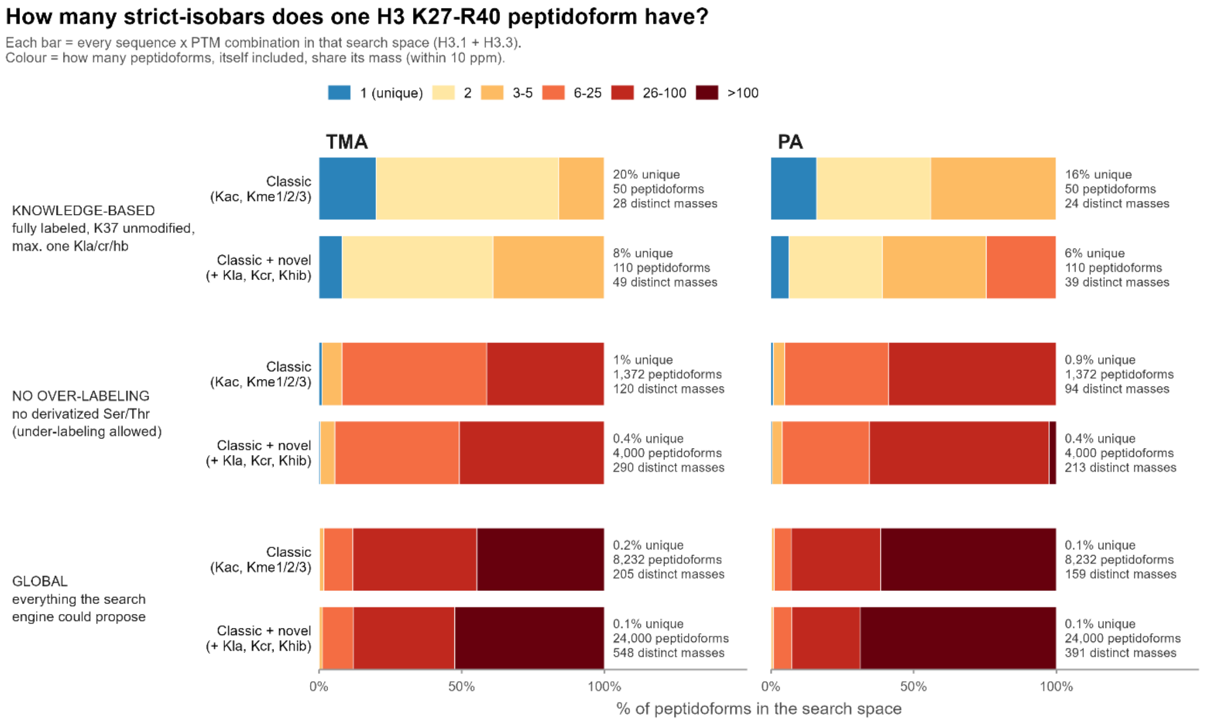


**Supplementary Figure 9: Theoretical calculation of the number of strictly isobaric peptidoforms corresponding to peptide H3 and H3.3 sequence stretch K27-R40, when using TMA as derivatizing agent and considering multiple lysine modifications**.

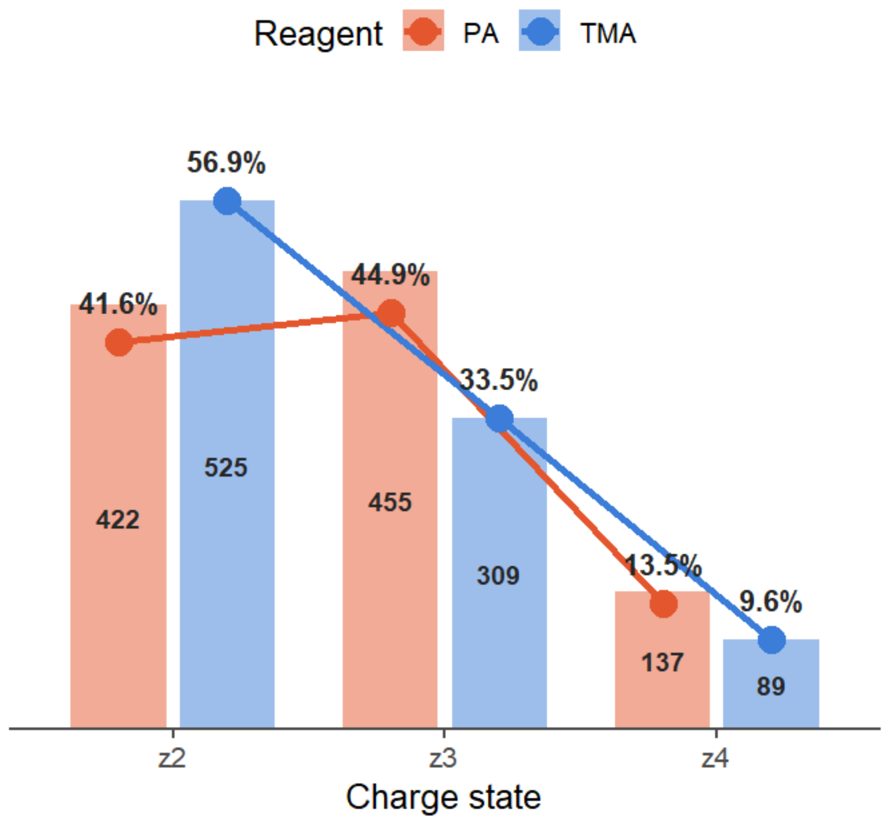


**Supplementary Figure 10: Comparison of the charge state distribution of histone peptides identified when using TMA or PA derivatization. Results obtained from the analysis of mouse testis histones.**

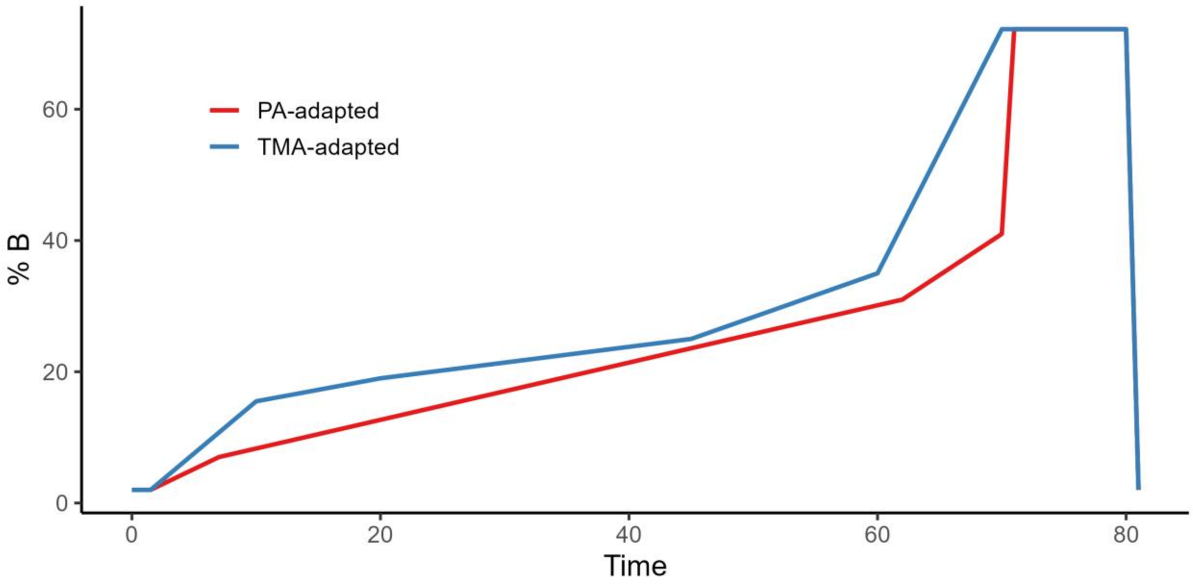


**Supplementary Figure 11**. **TMA-adapted liquid-chromatography gradient**. The gradient was adapted from previous gradient used for propionylated histones (PA-adapted) to account for increased hydrophobicity brought by the TMA moiety compared to propionyl. Mobile phase B is composed of ACN with 0.08 % (v/v) FA.

**Suppl. Table 1**. Acquisition parameters used on the Orbitrap Ascend Tribid MS instrument for analyses in Data-Dependent Acquisition (DDA) mode.

| | | |
|---|---|---|
| Global | Polarity | positive |
| | Fragmentation | HCD |
| | Range | 300 to 1300 m/z |
| Full MS Scans | Resolution | 120 000 |
| | Absolute AGC | 4e5 |
| | Max inj time | Auto |
| | Dynamic Exclusion | 10 s |
| | Isolation mode | Quadrupole |
| MS/MS Scans | Resolution | 30 000 |
| | Absolute AGC | 5e4 |
| | Max inj time | 59 ms |
| | NCE | 30 |
| | Isolation window | 1.5 m/z |

**Suppl. Table 2**. List of targeted Kacme1-containing peptides for validation using Parallel Reaction Monitoring (PRM)

| Mass [m/z] | CS [z] | Polarity | Start [min] | End [min] | NCE | Sequence |
|---|---|---|---|---|---|---|
| 422.748 | 2 | Positive | 11.09 | 17.09 | 27 | T[+84.057515]K[+56.026215]QTAR (light) |
| 427.7521 | 2 | Positive | 11.09 | 17.09 | 27 | T[+84.057515]K[+56.026215]QTAR (heavy) |
| 464.7844 | 2 | Positive | 37.21 | 43.21 | 27 | K[+140.08373]LPFQR (light) |
| 469.7885 | 2 | Positive | 37.21 | 43.21 | 27 | K[+140.08373]LPFQR (heavy) |
| 542.3115 | 2 | Positive | 12.37 | 18.37 | 27 | K[+140.08373]STGGK[+42.010565]APR (light) |
| 547.3156 | 2 | Positive | 12.37 | 18.37 | 27 | K[+140.08373]STGGK[+42.010565]APR (heavy) |
| 562.6736 | 3 | Positive | 12.28 | 18.28 | 27 | K[+112.088815]SAPATGGVK[+56.026215]K[+84.057515]PHR (light) |
| 563.335 | 2 | Positive | 19.89 | 25.89 | 27 | K[+140.08373]STGGK[+84.057515]APR (light) |
| 563.335 | 2 | Positive | 20.99 | 26.99 | 27 | K[+168.11503]STGGK[+56.026215]APR (light) |
| 566.0096 | 3 | Positive | 12.28 | 18.28 | 27 | K[+112.088815]SAPATGGVK[+56.026215]K[+84.057515]PHR (heavy) |
| 567.3454 | 3 | Positive | 12.23 | 18.23 | 27 | K[+126.104465]SAPATGGVK[+56.026215]K[+84.057515]PHR (light) |
| 568.3391 | 2 | Positive | 19.89 | 25.89 | 27 | K[+140.08373]STGGK[+84.057515]APR (heavy) |
| 568.3391 | 2 | Positive | 20.99 | 26.99 | 27 | K[+168.11503]STGGK[+56.026215]APR (heavy) |
| 570.6815 | 3 | Positive | 12.23 | 18.23 | 27 | K[+126.104465]SAPATGGVK[+56.026215]K[+84.057515]PHR (heavy) |
| 584.8561 | 2 | Positive | 28.54 | 34.54 | 27 | K[+140.08373]QLATK[+42.010565]AAR (light) |
| 589.8602 | 2 | Positive | 28.54 | 34.54 | 27 | K[+140.08373]QLATK[+42.010565]AAR (heavy) |
| 605.8795 | 2 | Positive | 40.66 | 46.66 | 27 | K[+168.11503]QLATK[+56.026215]AAR (light) |
| 610.8837 | 2 | Positive | 40.66 | 46.66 | 27 | K[+168.11503]QLATK[+56.026215]AAR (heavy) |
| 695.9007 | 2 | Positive | 38.6 | 44.6 | 27 | Y[+84.057515]QK[+56.026215]STELLIR (light) |

| | | | | | | |
|---|---|---|---|---|---|---|
| 700.9048 | 2 | Positive | 38.6 | 44.6 | 27 | Y[+84.057515]QK[+56.026215]STELLIR (heavy) |
| 715.8673 | 2 | Positive | 15.48 | 21.48 | 27 | D[+84.057515]AVTYTEHAK[+56.026215]R (light) |
| 720.8715 | 2 | Positive | 15.48 | 21.48 | 27 | D[+84.057515]AVTYTEHAK[+56.026215]R (heavy) |
| 733.4223 | 2 | Positive | 30.43 | 36.43 | 27 | D[+84.057515]NIQGITK[+56.026215]PAIR (light) |
| 738.3907 | 2 | Positive | 47.2 | 53.2 | 27 | E[+84.057515]IAQDFK[+56.026215]TDLR (light) |
| 738.4264 | 2 | Positive | 30.43 | 36.43 | 27 | D[+84.057515]NIQGITK[+56.026215]PAIR (heavy) |
| 743.3948 | 2 | Positive | 47.2 | 53.2 | 27 | E[+84.057515]IAQDFK[+56.026215]TDLR (heavy) |
| 762.9445 | 2 | Positive | 50.91 | 56.91 | 27 | V[+84.057515]TIMPK[+56.026215]DIQLAR (light) |
| 767.9487 | 2 | Positive | 50.91 | 56.91 | 27 | V[+84.057515]TIMPK[+56.026215]DIQLAR (heavy) |
| 832.0169 | 2 | Positive | 43.78 | 49.78 | 27 | G[+84.057515]K[+56.026215]GGK[+84.057515]GLG K[+84.057515]GGAK[+84.057515]R (light) |
| 837.021 | 2 | Positive | 43.78 | 49.78 | 27 | G[+84.057515]K[+56.026215]GGK[+84.057515]GLG K[+84.057515]GGAK[+84.057515]R (heavy) |
| 857.5041 | 2 | Positive | 21.4 | 27.4 | 27 | K[+140.08373]SAPATGGVK[+56.026215]K[+84.0575 15]PHR (light) |
| 862.5083 | 2 | Positive | 21.4 | 27.4 | 27 | K[+140.08373]SAPATGGVK[+56.026215]K[+84.0575 15]PHR (heavy) |
| 865.5016 | 2 | Positive | 20.48 | 26.48 | 27 | K[+140.08373]SAPSTGGVK[+56.026215]K[+84.0575 15]PHR (light) |
| 870.5057 | 2 | Positive | 20.48 | 26.48 | 27 | K[+140.08373]SAPSTGGVK[+56.026215]K[+84.0575 15]PHR (heavy) |
| 871.5198 | 2 | Positive | 28.91 | 34.91 | 27 | K[+168.11503]SAPATGGVK[+56.026215]K[+84.0575 15]PHR (light) |
| 871.5198 | 2 | Positive | 27.32 | 33.32 | 27 | K[+140.08373]SAPATGGVK[+84.057515]K[+84.0575 15]PHR (light) |
| 876.5239 | 2 | Positive | 28.91 | 34.91 | 27 | K[+168.11503]SAPATGGVK[+56.026215]K[+84.0575 15]PHR (heavy) |
| 876.5239 | 2 | Positive | 27.32 | 33.32 | 27 | K[+140.08373]SAPATGGVK[+84.057515]K[+84.0575 15]PHR (heavy) |
| 879.5173 | 2 | Positive | 26.32 | 32.32 | 27 | K[+140.08373]SAPSTGGVK[+84.057515]K[+84.0575 15]PHR (light) |
| 884.5214 | 2 | Positive | 26.32 | 32.32 | 27 | K[+140.08373]SAPSTGGVK[+84.057515]K[+84.0575 15]PHR (heavy) |
| 918.021 | 2 | Positive | 61.25 | 67.25 | 27 | K[+140.08373]TVTAM[+15.994915]DVVYALK[+84.05 7515]R (light) |
| 923.0251 | 2 | Positive | 61.25 | 67.25 | 27 | K[+140.08373]TVTAM[+15.994915]DVVYALK[+84.05 7515]R (heavy) |
